# Leveraging rDNA and rRNA dynamics to investigate the link between ecology and physiology in a freshwater microbial community

**DOI:** 10.64898/2026.02.24.707719

**Authors:** William R. Shoemaker, Martina Dal Bello, Jacopo Grilli

## Abstract

The dynamics of microbial communities in temporally varying environments are often mediated by the physiological responses of their members. However, physiological measurements remain difficult to obtain *in situ* from diverse communities in nature, limiting our understanding of microbial community dynamics in changing environments. Here, we investigate the physiological dynamics of a microbial community by applying a macroecological approach to a multi-year timeseries of paired ribosomal RNA (rRNA) and DNA (rDNA) measurements from a freshwater community, interpreting their relationship by developing a model of microbial physiology. Leveraging the existence of seasonal oscillations, we developed a data-driven minimal model that provided macroecological predictions and interpretations of rDNA, rRNA, and their relationship. Using this model, we examined two macroecological patterns with physiological interpretations: 1) time averaged rRNA:rDNA as a measure of a community member’s steady-state and 2) the temporal ordering of rRNA fluctuations relative to rDNA. We found that rRNA:rDNA increased with the number of 16S rRNA operon copies, which in the context of our model suggests a superlinear effect of copy number. Temporal fluctuations of rRNA tended to *precede* those of rDNA, consistent with the view that increases in ribosomal content can lead to future biomass. We then identified environmental variables where rRNA and rRNA:rDNA displayed greater sensitivity than rDNA, reasoning that rRNA:rDNA, as a reflection of the growth rate, may respond more strongly to environmental change. The ratio rRNA:rDNA provided greater sensitivity than rDNA for the majority of the aquatic environmental variables tested, particularly those reflecting the degree of eutrophy vs. oligotrophy. The results of this work indicate that the ratio rRNA:rDNA may be underutilized as a measure of the physiological state of natural communities.

## Introduction

The physiological response of microorganisms to changing environmental conditions shapes the composition and stability of the communities they inhabit [1–4]. Cellular physiology therefore provides a mechanistic grounding for observed ecological dynamics [1, 3, 5, 6]. Because cells contain myriad physiological features, it is necessary to focus on those that exert the strongest control over growth. Microbiological investigations have pared down this physiological complexity into a set of empirical relationships known as “growth laws,” a prominent example being that the growth rate is directly related to the fraction of mass that is allocated to ribosomes by the cell [7–10]. Such investigations have primarily been performed for *Escherichia coli* under controlled experimental conditions, though recent studies demonstrate their applicability to phylogenetically diverse bacterial species [11] and models of microbial community dynamics [12]. However, while physiology plays a clear role in the growth dynamics of microbial communities, obtaining physiological measures in natural microbial communities has proven to be a difficult task.

Due to the inherent difficulty of directly assessing the physiological state of diverse *in situ* communities, sequence-based proxies are often measured. A frequently used approach is to sequence transcripts of the 16S rRNA gene (i.e., rRNA) in addition to DNA (i.e., rDNA) [13–26]. The ratio of these two measures, rRNA:rDNA, is often interpreted as a proxy for the metabolic activity of community members [16, 17, 20, 27–32]. This activity-based interpretation ultimately rests on the assumption that rRNA:rDNA reflects a community member’s ribosomal content, which confers the potential for growth [19, 23]. While alternative approaches exist for assessing growth in natural communities, they are limited in that they are difficult to apply to high richness communities (e.g., stable isotope probing; [33]), are restricted to certain phylogenetic groups (e.g., spore-formation in Bacillota as a conserved life-history strategy; [34]), or are reflective of DNA replication rather than cellular physiology (e.g., relationship between coverage at a nucleotide site and distance to the origin of replication; [35, 36]). Instead, ribosomal transcript-based approaches have the advantage of leveraging a universal feature of microbial cells and often being proportional to cellular ribosome content, which itself can be proportional to the rate of growth when the latter is constant with respect to time [7]. However, we currently lack a quantitative understanding of the typical fluctuations and dynamics of rRNA relative to those of rDNA as well as whether their joint relationship reflects microbial physiology.

Identifying when and how rRNA differs from rDNA requires a quantitative, flexible framework. Within microbial ecology, macroecological principles have allowed researchers to characterize what one should expect in a typical community [37–44], spurring the development of novel mathematical models designed to capture empirical patterns with minimal fine-tuning [45–52]. For example, a data-driven mathematical model of density-dependent growth with environmental noise, the Stochastic Logistic Model of growth (SLM), has been successfully applied to a diverse array of macroecological patterns in both natural [53–55] and experimental communities [56], providing a foundation for predicting and interpreting novel macroecological patterns [57–62]. Given the demonstrated success of macroecology in explaining and predicting patterns in rDNA, this same data-driven approach can be extended to rRNA, providing a principled way to determine how rRNA differs from rDNA as well as how they are jointly related. Because such models are constrained by empirical observation rather than fine-tuned, this approach offers a route to extracting physiological insight from the relationship between rRNA and rDNA (e.g., the ratio rRNA:rDNA).

Proxies of cellular physiology have the potential to inform our understanding of community-level responses to changing environmental conditions. Given that they are thought to track a community member’s physiological state, rRNA and rRNA:rDNA may display greater sensitivity to environmental change, registering shifts in growth-relevant conditions earlier, or more strongly, than corresponding changes in rDNA. Aquatic bacterioplankton communities provide an ideal system for assessing the environmental dependency of physiological measures, as seasonality is expected to generate a periodic signature on abundance dynamics [63–66], a form of environmental variation that may impact cellular ribosomal content. For example, aquatic communities routinely transition between oligotrophic (i.e., nutrient-poor) and eutrophic (i.e., nutrient-rich) conditions [66, 67], driving shifts in the relative dominance of taxa that differ systematically in growth rate and, by extension, ribosome content [7, 68, 69]. Similarly, rRNA may be highly sensitive to unambiguously abiotic factors such as temperature, as it shapes all three processes of the central dogma as well as ribosome assembly [70], with physiology exhibiting consistent temperature-dependency across phylogenetically diverse species [71, 72] that consequently provide a plausible mechanistic link between environmental variation and the community-level dynamics documented in natural microbial communities [73–79]. However, the sensitivity of rRNA as a measure of physiology relative to rDNA as a measure of population abundance has yet to be examined in natural communities.

In this study, we investigated the dynamics of rDNA and rRNA in a freshwater community over a multi-year timescale [15]. The dynamics of the system occurred over two characteristic timescales: short-term fluctuations and long-term periodic oscillations. This separation of timescales allowed us to model oscillatory dynamics as a gamma distribution, the stationary distribution of the SLM, with a time-varying mean. Fluctuations and dynamics of rDNA and rRNA were highly consistent across community members, displaying similar oscillation timescales and amplitudes, reflecting their physiological coupling. The rRNA:rDNA ratio as a proxy for ribosomal mass fraction was consistent with predictions of average growth *across* community members and rRNA fluctuations preceded those of rDNA, both of which could be recapitulated by a physiological model of population dynamics. Finally, we investigated the magnitude of change in predicted rRNA:rDNA in response to environmental variation (i.e., sensitivity), reflecting a shared physiological response to environmental change. The results of this study provide insight into the typical response, variation in response, and degree of information provided by rRNA as a proxy for microbial physiology in an aquatic environment.

## Results

### rRNA and rDNA display concordant and divergent seasonal dynamics in an aquatic microbial community

In this work, we considered a freshwater microbial timeseries, weekly sampled for *≈*2.5 years [15]. In this study, rRNA was measured by reverse transcribing environmental RNA into complementary DNA, the 16S rRNA region of which was then amplified and sequenced alongside paired environmental DNA (Fig. 1a). Figure 1b shows that read counts of both rDNA and rRNA of individual community members displayed oscillations as a form of seasonality. These oscillations occurred over an extended timescale, ranging from the order of months to a year. The most prominent example of such dynamics was the dominant community member, belonging to the family of photosynthetic cyanobacteria Nostocaceae, with seasonal variation in relative abundance of over four orders of magnitude. Given the diurnal cycle of the environment, the timescale of cellular division is expected to be much smaller than these observed oscillations. Therefore, we modeled abundance dynamics as a combination of short-term fluctuations and long-term deterministic seasonal trends to facilitate rRNA and rDNA comparison. These long-term dynamics were incorporated into the Stochastic Logistic Model (SLM) as a time-dependent carrying capacity that oscillates over environmental timescale τ ^env^ (Fig. 1c). Macroecological measures of abundance supported this choice of model, as distributions of abundances over time (the Abundance Fluctuation Distribution) resembled the stationary distribution of the SLM (i.e., the gamma), and the sampling form of the gamma recapitulated the relationship between ASV abundance and occupancy (i.e., fraction of samples where the ASV was observed; Fig. S1a,b, S3) for both rRNA and rDNA. In addition, the exponent of the relationship between the mean and variance of abundance (i.e., Taylor’s Law) for rRNA and rDNA was nearly identical (Fig. S1c).

These results suggest that the same ecological model can be applied to rRNA and rDNA, from which the exponent of Taylor’s Law implied a similar dependence between the carrying capacity and the strength of environmental noise for both rDNA and rRNA. Per-community member measures including mean relative abundance, occupancy, and the correlation between pairs of community members also had similar values for rDNA and rRNA (Fig. S2). These results indicate that rRNA and rDNA are virtually indistinguishable in terms of general macroecological patterns. We note that these patterns do not consider time, suggesting that rRNA and rDNA are effectively indistinguishable for time-agnostic macroecological patterns.

Accurate inference of model parameters required both an appropriate data transformation and a statistical inference procedure. Microbial 16S rRNA amplicon data are fundamentally compositional in nature, meaning that the data can only contain relative information, as the sum of read counts across community members must be equal to the total number of reads in that sample. This constraint means that common transformations such as taking the relative abundance of read counts can cause artifacts to appear in the data, where an *increase* in the relative abundance of one community member can make it appear as though another community member is *decreasing* in abundance, when its true abundance remains unchanged. The issue of compositionality is clearly pertinent for assessing the validity of oscillatory dynamics, and transformations that explicitly account for data compositionality have been advocated [80–82]. Therefore, we inferred model parameters via numerical maximization of the gamma distribution (i.e., the stationary distribution of the SLM) with oscillating mean (i.e., carrying capacity; Eq. 9) using CLR-transformed read counts for community members that were present in every sample. Simulations confirmed that this approach was appropriate for accurate inference (Supporting Information; Figs. S4, S5, S6) as did additional analyses of potential data collection confounders (Supporting Information; Figs. S7, S8).

**Figure 1.**
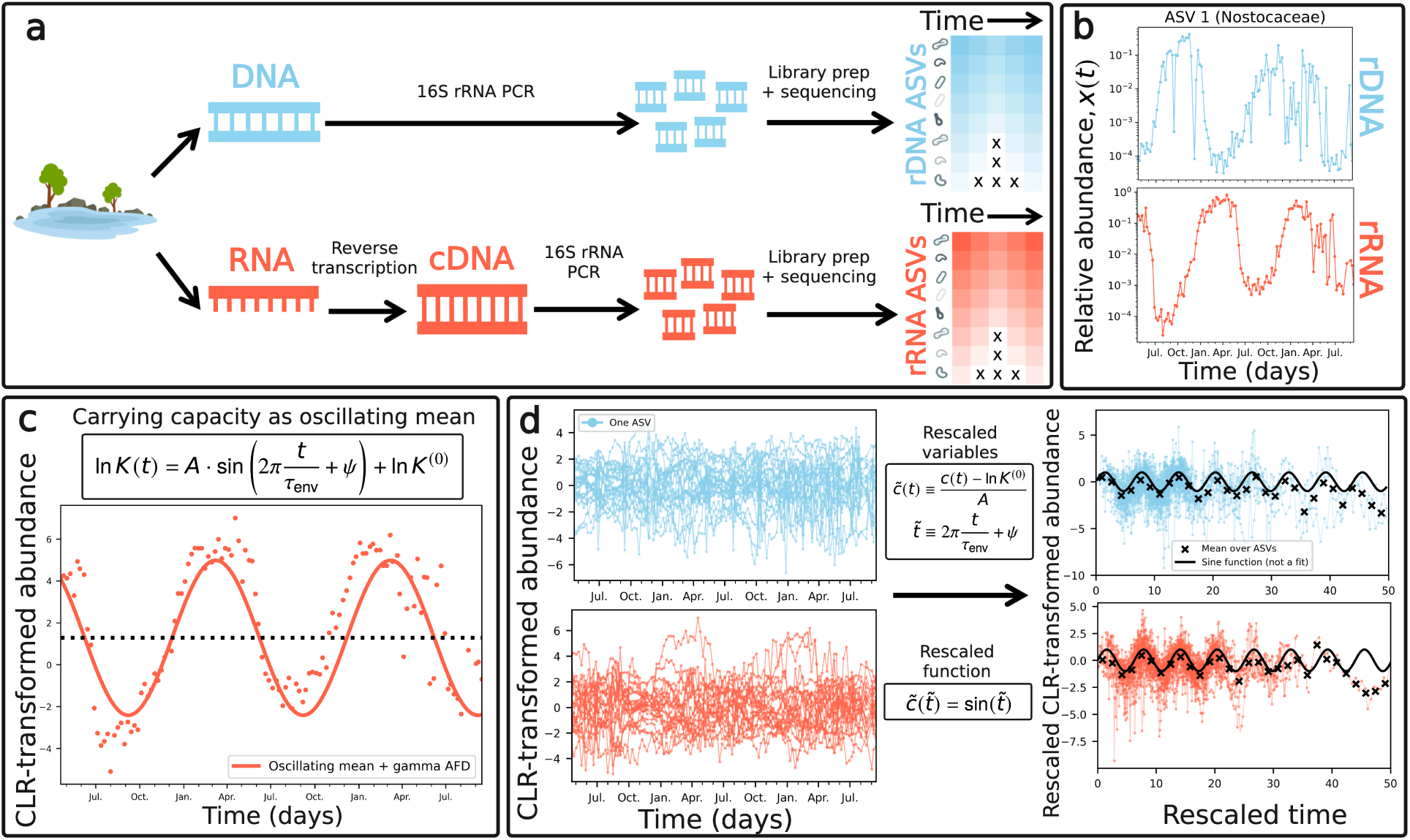
Timeseries of rDNA and rRNA reveal shared seasonal dynamics across community members in a freshwater lake. **a)** A prior study sequenced both the rDNA and the reverse-transcribed rRNA of the 16S rRNA gene in a freshwater lake *∼* weekly over the span of 28 months. This dataset provided the means to examine the dynamics of rDNA as a reflection of biomass and rRNA as a reflection of both biomass and ribosome content across an assemblage of microbial taxa. **b)** When relative abundance was plotted over time, many community members exhibited clear oscillatory dynamics for both rDNA and rRNA. **c)** Observed oscillatory dynamics were incorporated into the stationary form of the Stochastic Logistic Model by defining the logarithm of the carrying capacity as a sinusoidal function. **d)** These carrying capacity parameters varied across community members for both rDNA and rRNA timeseries, making it difficult to assess whether taxa with different parameters exhibited similar dynamics. By rescaling abundances and time for each community member, we identified similar population dynamics, known as a data collapse.

We found high agreement between the fitted seasonality model and the CLR-transformed abundance data for both rRNA and rDNA (Figs. S9, S10). Our model was also used to predict the correlation between the present and future abundance of a community member (Supporting Information), where we found reasonable agreement between empirical autocorrelation and the predictions of the oscillatory model (Figs. S11, S12). Different oscillation parameters reflect considerable variation across community members. Such variation became apparent when abundance trajectories of all community members were simultaneously plotted, as the data did not collapse on a single curve (Fig. 1d). Once abundances were appropriately rescaled using our inferred parameters, we found that all community members fell on a single sinusoidal curve, a phenomenon known as data collapse [83, 84]. The ability to take quantitatively dissimilar variables and reveal their qualitatively similar form is not trivial, as simulations confirmed that the existence of the data collapse was not an artifact of statistical inference (Fig. 1d); rather, it can be interpreted as a reflection of the underlying shared dynamics of community members (Fig. S15).

**Figure 2.**
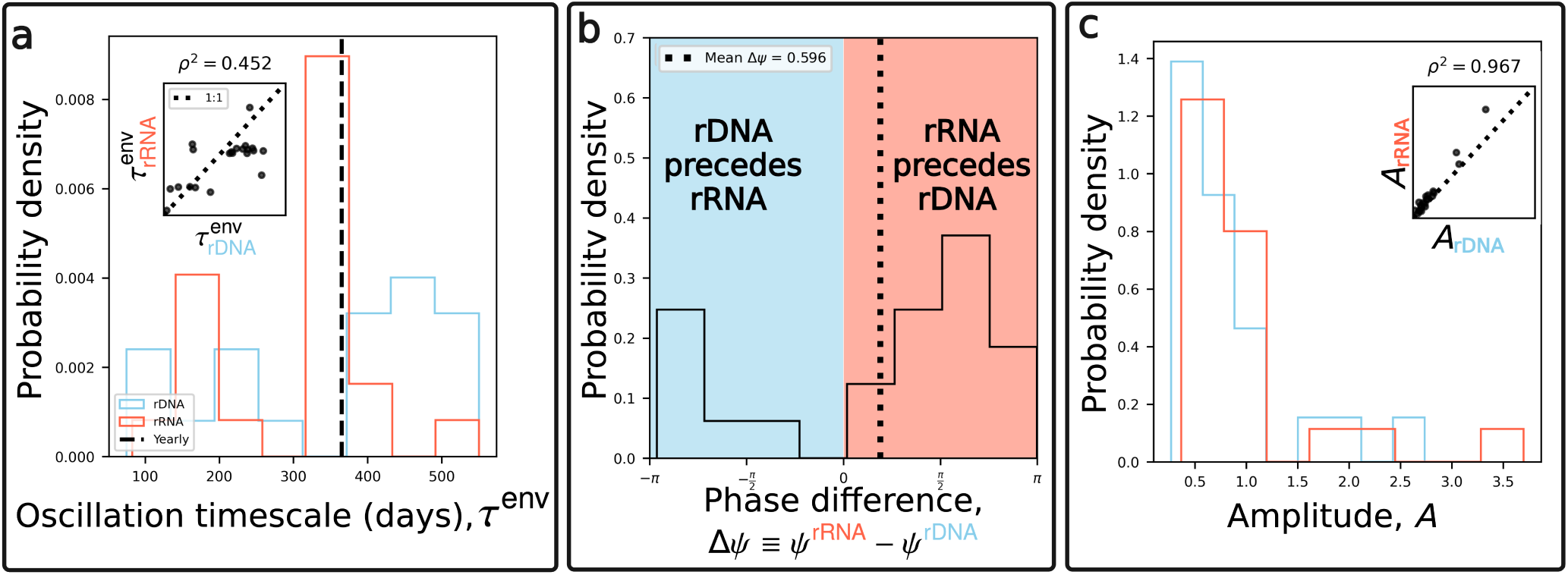
rDNA and rRNA oscillations are strongly coupled via governing parameters with temporal ordering between them. Variation in oscillatory parameters between rRNA and rDNA was assessed. **a)** Distributions of oscillation timescales τ ^env^ largely overlapped for both rDNA and rRNA, and timescales were significantly correlated across community members (inset figure). Both distributions exhibited strong peaks corresponding to yearly timescales (365 d), with smaller peaks corresponding to semi-yearly (*∼* 180 d) and seasonal (*∼* 90 days). **b)** The difference in oscillatory phase parameters was examined as a measure of the order in which rRNA and rDNA oscillated, reflecting whether a change in rRNA abundance followed a change in rDNA or vice versa. While the bulk of the distribution was centered near zero, reflecting close coupling between rRNA and rDNA oscillations, a wrapping-invariant test of phases on the unit circle confirmed that this distribution was non-uniform across ASVs, indicating a modest but systematic tendency for one measure to precede the other (Fig. S16). **c)** The distribution of oscillation amplitudes largely overlapped for rRNA and rDNA and exhibited a strong correlation (inset figure), reflecting the strength of the coupling between rDNA and rRNA. We note ASV 1, the sole phototroph in the community, as a clear outlier.

### rRNA and rDNA displayed similar oscillatory dynamics across community members

Systematic deviations between rRNA and rDNA that hold across community members have the potential to reveal the physiological state of a dynamic community. Using the inferred oscillatory parameters, we assessed the similarity between rDNA and rRNA. First, by plotting the distribution of τ ^env^, we found similarities between rDNA and rRNA. Both rDNA and rRNA displayed two characteristic oscillation timescales, approximately corresponding to oscillations occurring annually and semi-annually (Fig. 2a). We note that the width of the distribution around the yearly timescale was considerably narrower for rRNA, potentially due to its greater sensitivity to changing environmental conditions. In addition, oscillation timescales between rDNA and rRNA were significantly correlated across ASVs, likely reflecting the physiological coupling of the two variables.

We then asked whether increases in rRNA preceded those of rDNA or vice versa. If an increase in rRNA reflects the buildup of ribosomes that precedes biomass formation, then rRNA increases should systematically lead rDNA increases. Therefore, any systematic tendency in the order of oscillations can provide information about the applicability of rRNA as a reflection of the physiological state of community members. In terms of our inferred parameters, we investigated the sign of the difference between the phases, Δψ *≡* ψ^rRNA^ *−* ψ^rDNA^. The distribution of phase differences across community members deviated from zero, suggesting that rRNA and rDNA oscillations were *ordered* across ASVs (Fig. 2b). While we plotted Δψ as a distribution for visualization purposes, statistical tests of phases on a one-dimensional axis are susceptible to phase wrapping. Therefore, we performed a wrapping-invariant test, finding that ASVs were non-uniformly distributed on the unit circle (Fig. S16; R = 0.484, P = 0.006; Supporting Information).

**Figure 3.**
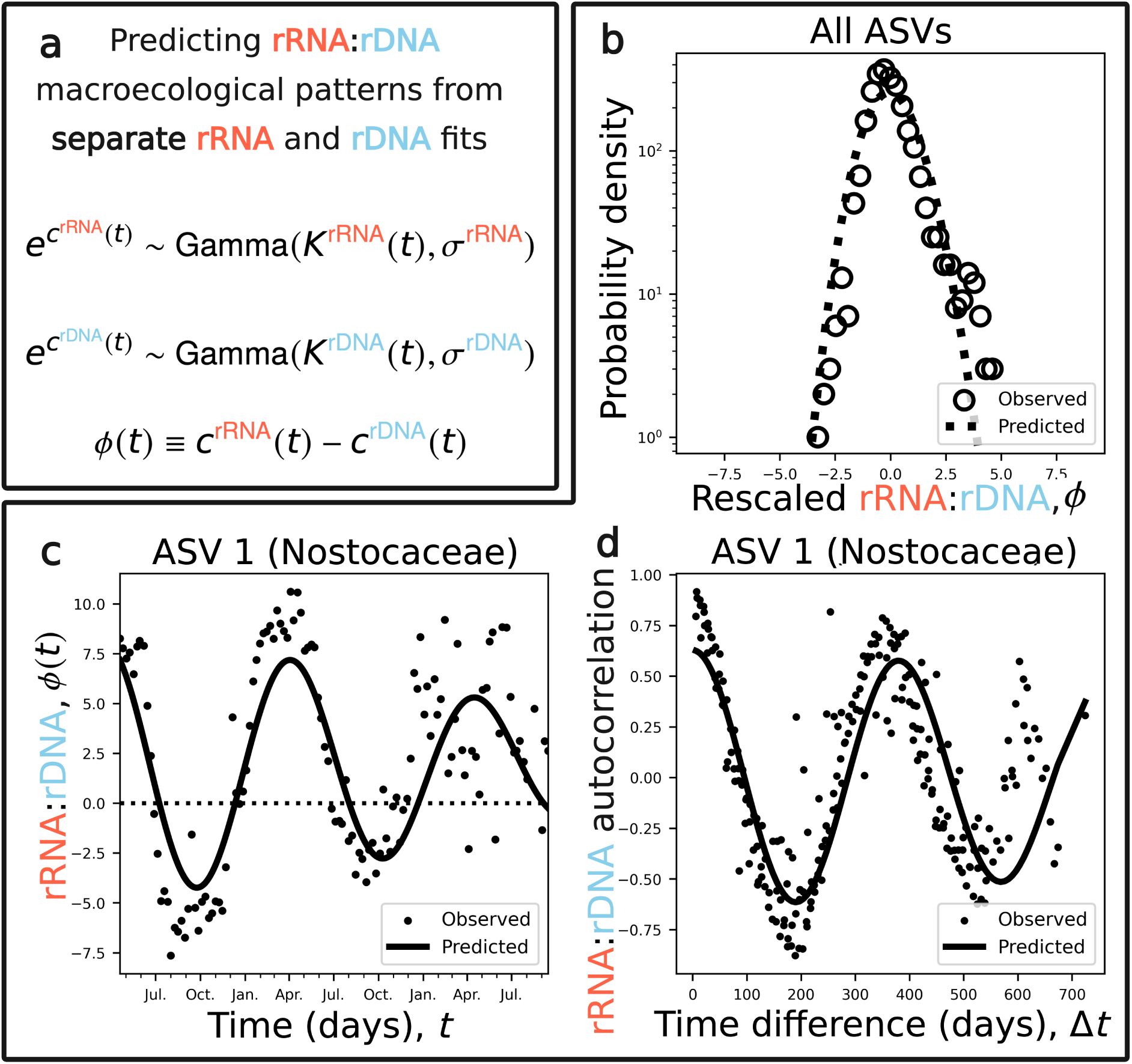
Predicting macroecological properties of rRNA:rDNA. **a)** Given that the abundances of rRNA and rDNA are each well-described as independent gamma random variables with time-varying means, one can derive and test macroecological predictions of rRNA:rDNA without performing additional fitting. **b)** The distribution of rRNA:rDNA as the ratio of two gamma-distributed random variables can be derived and sufficiently captures the empirical distribution over all ASVs using separate rRNA and rDNA fits. The temporal dynamics of rRNA:rDNA are also well-described by our model for individual ASVs, both for **c)** the change in rRNA:rDNA over time and **d)** its temporal autocorrelation.

Finally, we investigated the amplitude as a way to identify patterns in the strength of oscillations across community members. The distributions of both rDNA and rRNA were highly similar, with both datasets displaying peaks at *∼* 0.5 and amplitude values being highly correlated (Fig. 2c). However, both distributions contained a clear outlier that was not present in the distributions of timescales or phase differences. This outlier was identified as the most abundant community member, a phototroph in the family Nostocaceae. We briefly note that there was no clear relationship between the pairwise phylogenetic distance between community members and the difference in their oscillation parameters (Fig. S14). This observation can be interpreted as variation in the carrying capacity not being related to shared evolutionary history captured by the 16S rRNA gene.

### Predicting rRNA:rDNA macroecology

In the previous section, we found that rRNA and rDNA often exhibited similar oscillation timescales and amplitudes, suggesting that each measure was effectively capturing the same underlying abundance dynamics. The *ratio* of these two variables is often interpreted as a measure of cellular activity (rRNA:rDNA; e.g., [17, 19]), which here we specify as being reflective of the ribosomal content of a community member. Therefore, to determine whether rRNA differs from rDNA, we tested whether our previously introduced model of rRNA and rDNA as statistically independent random variables (i.e., no joint relationship between rRNA and rDNA) with time-varying means could capture macroecological patterns of rRNA:rDNA (ϕ*_i_*(t); Fig. 3a). Using the observation that rRNA and rDNA are well-described by a gamma distribution (Fig. S1), we derived a predicted distribution that succeeded in capturing the empirical distribution of rRNA:rDNA (Fig. 3b). This success extended to temporal macroecological patterns of individual ASVs, as the predicted change in rRNA:rDNA over time and its temporal autocorrelation showed reasonable agreement with the data (Fig. 3c,d; S17, S18).

### rRNA:rDNA as a reflection of microbial physiology across and within community members

The ratio rRNA:rDNA is often interpreted as a measure of the ribosomal content of a microbial species, a key determinant of growth in experimental settings. Thus, we asked whether the relationship between rRNA and rDNA reflected predictions from microbial physiology both *across* community members averaged over time and *within* a given community member over time. To answer these questions, we developed a minimal mathematical model that explicitly couples rRNA and rDNA dynamics via cellular growth while maintaining both logistic growth and the same stationary distribution as the SLM (i.e., gamma). Environmental noise, a key ingredient of the SLM, was incorporated by treating the rates of cellular growth, ribosome assembly, and ribosome degradation as stochastic processes (Fig. 4; Supporting Information).

We first focused on how rRNA:rDNA varied *across* community members, examining whether average rRNA:rDNA was related to proxies of microbial growth. Specifically, for bacteria, the growth rate has been found to be a function of the number of 16S rRNA operon copies housed in the genome [68, 85–89], a relationship that has been leveraged to investigate the seasonal dynamics of marine microbial communities [78, 90]. If rRNA:rDNA reflects ribosomal content, and, thus, growth, we would expect rRNA:rDNA to increase with operon copy number, a result predicted by the steady-state solution of our model (Fig. 4b). By averaging ϕ*_i_*(t) with respect to time for each community member, we found evidence of said relationship (α = 1.65, t = 4.24, P = 0.00819; Fig. 4b). While it is difficult to interpret the value of this slope given the moderate number of ASVs and compositional nature of the data, in the context of our model a positive slope represents a *superlinear* gene dosage effect. An equivalent relationship was found using the inferred parameter that reflected the time-average over a single oscillatory cycle in our gamma model (i.e., K^(0)^; Fig. S19; Supporting Information).

Moving from steady-state to temporal dynamics, we sought to provide a mechanistic understanding of the systematic tendency for increases in rRNA to *precede* those of rDNA. In the previous section, we investigated the temporal ordering at a single frequency. To provide greater rigor and obtain a physiological interpretation from our model, we examined phase delay using the Fourier transform of the cross-covariance function (i.e., cross-power spectral density; Fig. 4c; Supporting Information). Here, rRNA *precedes* rDNA (i.e., phase delay is positive) when the ratio of ribosomal assembly vs. degradation exceeds that of cellular vs. ribosomal noise, an inequality we plotted as a function of ribosomal noise strength increasing while all other parameters remain fixed (Fig. 4c). A positive phase lag was observed for the majority of ASVs, a result that was statistically significant at the global level when compared to a phase-randomized null distribution (Fig. 4c, S20).

**Figure 4.**
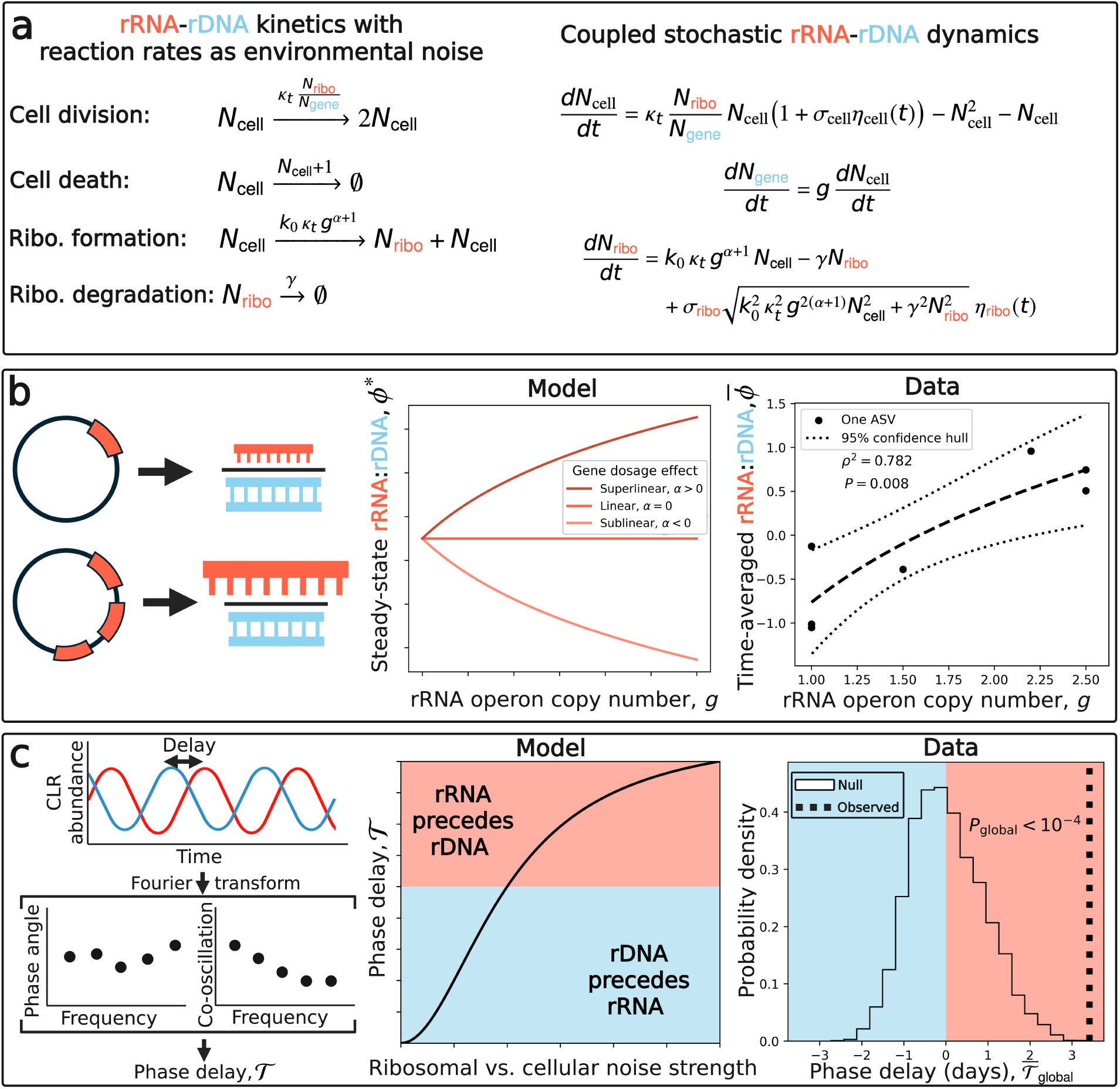
Testing physiological predictions of rRNA:rDNA across and within community members. The ratio between rRNA and rDNA relative abundances has frequently been interpreted in microbial ecology as a proxy for a community member’s ribosome content and, thus, a reflection of growth. **a)** We formalized this interpretation as a mechanistic model, allowing reaction rates to stochastically vary as a form of environmental noise. **b)** At steady-state, the model predicted a positive relationship between 16S operon copy number and rRNA:rDNA due to superlinear gene dosage effects, a relationship that was empirically observed across ASVs. **c)** The temporal ordering of the model was derived as a phase lag, wherein changes in abundance of rRNA preceded those of rDNA when ribosomal noise became sufficiently large. This positive phase lag was statistically significant when aggregating across ASVs, wherein a null distribution was obtained by randomizing phases (i.e., randomized temporal ordering).

### Assessing global sensitivity of rRNA and rRNA:rDNA to oscillating environmental variables

Measurements of rRNA are often reported to have greater sensitivity to environmental variables than those of rDNA. If rRNA:rDNA reflects growth rate, we would expect it to respond more strongly than rDNA to temporal changes in environmental variables. To investigate this claim, we leveraged paired measurements of bulk, oscillating environmental variables (Fig. 5a). Due to the non-linearity of our measurements, we identified and fit an appropriate Generalized Additive Model (GAM) to each ASV with environmental variables as predictors and rRNA, rDNA, and rRNA:rDNA as our three response variables (Supporting Information). Therefore, *sensitivity* was defined as the derivative of the relationship between *predicted* CLR-transformed abundance and a given environmental variable (Fig. 5b).

Through this approach, we tested whether rRNA and/or rRNA:rDNA displayed greater sensitivity than rDNA for nine bulk environmental variables. The ratio rRNA:rDNA displayed significantly greater sensitivity for the majority of environmental variables (6/9; Fig. 5c), suggesting that paired measurements of rRNA and rDNA may serve as appropriate measurements for investigating how microbial communities respond to changing environmental conditions. Notably, three of these six variables, dissolved O_2_, Secchi depth, and dissolved organic carbon (DOC), can be collectively interpreted as a single environmental axis that distinguishes oligotrophic from eutrophic conditions, suggesting that sensitivity reflects the ratio’s responsiveness to the trophic state of the community.

In contrast, rRNA alone only had greater sensitivity than rDNA for Secchi depth and actually displayed *lower* sensitivity than rDNA for salinity, specific conductivity, and total phosphorus. While no individual ASV disproportionately contributed to inferred sensitivity (Supporting Information), we note that the correlation in sensitivities between ASV pairs weakly decayed with increasing phylogenetic distance for both rRNA and rDNA (Fig. S21), potentially reflecting a conserved evolutionary physiological response.

**Figure 5.**
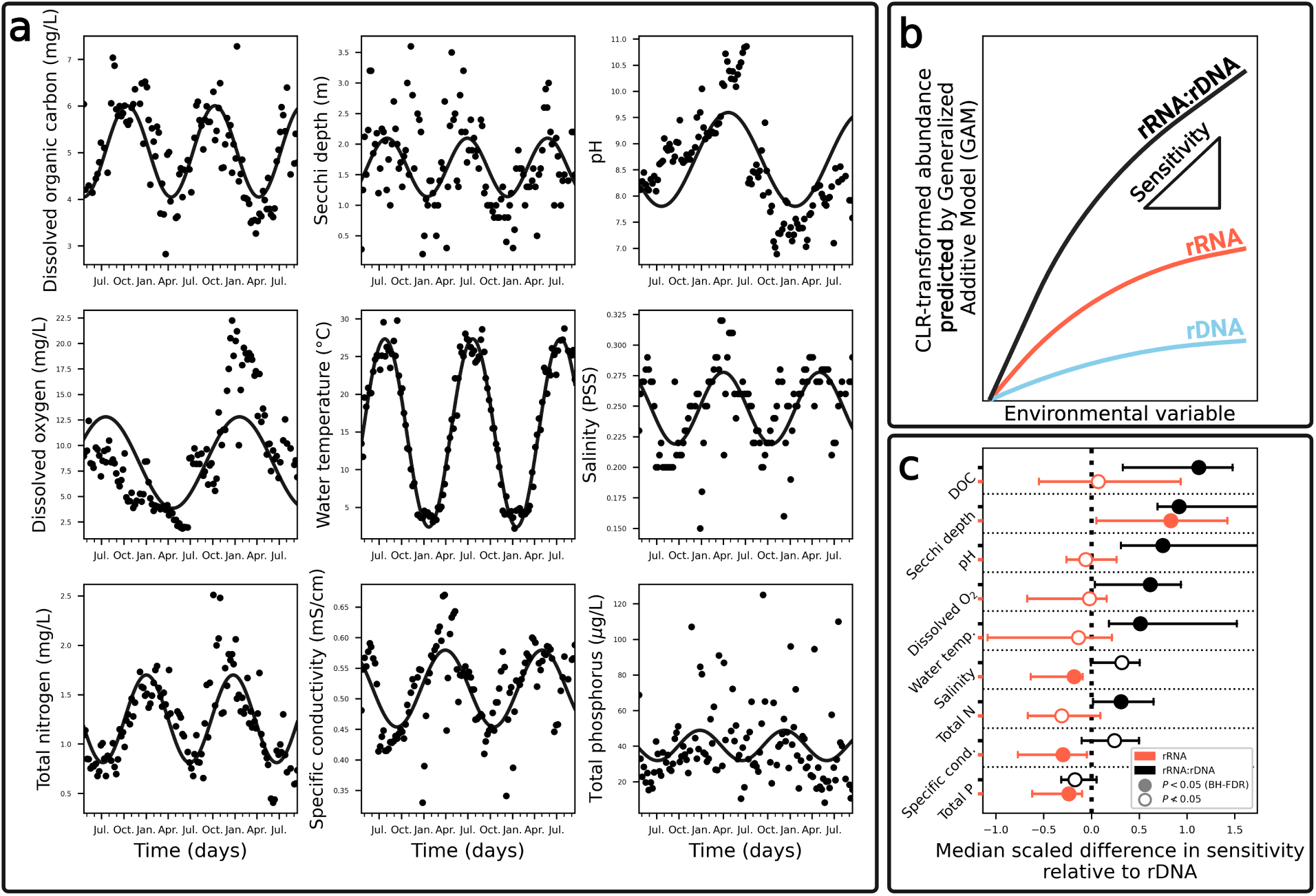
rRNA and rRNA:rDNA provide greater sensitivity than rDNA to select environmental factors. **a)** Similar to ASVs, environmental variables displayed oscillations of varying timescales. **b)** The sensitivity of rRNA and rRNA:rDNA relative to rDNA was assessed using Generalized Additive Models (GAMs), where the gradient of the *predicted* increase in abundance was estimated for each environmental variable for each ASV (Supporting Information). **c)** Using this approach, we identified the environmental variables where measuring rRNA provided an advantage over rDNA alone. The false-discovery rate was controlled for using the Benjamini–Hochberg procedure.

## Discussion

The dynamics of 16S rRNA and rDNA were highly similar across community members, displaying oscillations with similar amplitudes and timescales. By combining these qualitatively similar dynamics with fluctuations similar to those previously observed in non-oscillating communities, we developed a minimal model to investigate the variability of rRNA:rDNA, a ratio often examined in microbial ecology and environmental microbiology as proxy for growth rate. The distribution of rRNA:rDNA as well as the temporal dynamics and autocorrelation of the ratio could be predicted, providing a flexible, quantitative null model for investigating rRNA:rDNA.

Leveraging the interpretable nature of rRNA, rDNA, and rRNA:rDNA macroecological patterns, we identified steady-state and temporal patterns that held across community members, patterns that were successfully interpreted through a physiological minimal model of rRNA and rDNA dynamics. Starting with steady-state predictions, the observed positive relationship between 16S operon copy number and rRNA:rDNA *across* community members is driven by copy number having a superlinear effect on ribosome formation in our model. Assuming that rRNA:rDNA reflects the potential *in situ* growth rate, the existence of this relationship is consistent with prior experimental work confirming a positive relationship between copy number and growth rate across phylogenetically diverse bacteria [68, 85–90]. This relationship has been interpreted as reflecting an underlying trade-off between slow-growing oligotrophs and fast-growing copiotrophs, a microbial life-history strategy [68, 69], and reflects predictions from computational models of cellular physiology where the ratio of rRNA transcripts to biomass increased with the rRNA operon copy number [91]. We note that while members of the freshwater community we examined are unlikely to be maintaining a maximum growth rate given observed environmental oscillations, it is possible that time-averaged rRNA:rDNA is reflective of the time-averaged rate of growth.

Turning to temporal patterns, shifts in rRNA abundance consistently preceded those of rDNA, a result that was statistically significant at the community-level. This sequence of events is biologically intuitive, as increased ribosomal content may be necessary for the generation of additional protein mass, which comprises the bulk of the cell. To provide physiological context for when this ordering does and does not occur, we derived the phase lag from our model, finding that the observed temporal ordering reflected the ratio of the rates of ribosomal assembly vs. degradation exceeding that of cellular vs. ribosomal noise. We note that this ordering is reversed (i.e., rDNA *precedes* rRNA) when ribosomal noise is absent, meaning that accounting for ribosomal fluctuations is likely necessary for investigating the physiological dynamics of microbial communities. We also note that the sampling timescale is a limiting factor in our ability to extract physiological signals from these paired measurements. As a heuristic explanation, the taxonomic family of our dominant phototroph has a minimum division time (i.e., inverse of maximum growth rate) at optimal temperature of *∼* 0.33-5d under arguably ideal experimental conditions [92], whereas the median time between sampling events in our study was one week. Similar to conclusions drawn in recent investigations of the limits of inference in stochastic microbial timeseries [44, 93], we argue that stronger signals could likely be obtained when sampling resolution relative to the timescale of growth is explicitly taken into account.

The choice to measure rRNA is often justified by appealing to the transient nature of transcripts in the environment relative to DNA [23, 94]. We tested this claim by examining the sensitivity of rRNA or rRNA:rDNA vs. rDNA, defined as the absolute value of the derivative of predicted abundance with respect to a given environmental variable. Broadly, rRNA:rDNA displayed greater sensitivity to the majority of environmental variables because it leveraged scenarios when the derivative of rRNA and rDNA had different signs (i.e., + vs. -). It is beyond the scope of this work to propose mechanistic models for each environmental variable. However, it is worth briefly establishing broader connections between these results and the environmental microbiology literature. The significance of dissolved O_2_, Secchi depth, and DOC can all be interpreted as indexing a single environmental axis that captures oligotrophic vs. eutrophic conditions. In addition, prior studies found that rDNA exhibited a stronger response than rRNA to low dissolved O_2_concentrations [67], explaining the significant sensitivity for rRNA:rDNA. The significant response of rRNA:rDNA to pH could be attributed to either enzyme kinetics or membrane function [95], while the nitrogen response may be driven by regulation of nitrogen cycle pathways [96]. The temperature outcome is of particular interest given that its impact on growth across microbial lineages is comparatively well-established, both at the level of cells [70, 97–99] and communities [78, 79, 100, 101]. In contrast, rRNA displayed significantly *lower* sensitivity relative to rDNA to salinity, specific conductivity, and total phosphorus. The reduced sensitivity of rRNA to salinity is notable given recent efforts demonstrating that the environmental variable contributes to growth in aquatic microbial communities [90]. We note, however, that this effort focused on marine systems while this examines a freshwater community.

While 16S rRNA transcripts have been used in one form or another in microbial ecology for > 30 years [13], our results suggest that the tool may remain underutilized for physiological investigations. The timescale of sampling relative to that of cellular division is essential, as previously discussed, but measuring rRNA in response to perturbations and obtaining absolute abundances are additional considerations.

Regarding perturbations, we argue that there is sufficient temporal variation in our aquatic environment, occurring over slow (i.e., seasonal) timescales, to elicit detectable physiological responses. While sampling resolution is a limitation, this variation likely extends to unmeasured, fast (i.e., diurnal) timescales as well. Indeed, perturbation-based studies of soils with short measurement timescales (e.g., hours) found detectable rRNA responses [102–104]. In contrast, systematic perturbations and subsequent temporal measurements have not typically been applied in experimental studies that found 16S rRNA transcripts to be uninformative about growth (e.g., [105]). Absolute abundances have been obtained in prior studies via spike-ins [106], though temporal dynamics were not assessed, nor were environmental perturbations applied. Alternative rRNA-based approaches with greater physiological sensitivity may be leveraged alongside these experimental considerations. For example, the rRNA operon is initially transcribed into a single precursor known as pre-rRNA [107]. The concentration of pre-rRNA has been shown to be sensitive to the physiological state of the cell, and while it has a far lower concentration than that of ribosome-bound 16S rRNA, it can be sampled through modified barcoding approaches [108–110].

## Materials and methods

### Data acquisition

Raw 16S barcode data were acquired from a previously published study [15] and reprocessed using dada2 v1.26 [111] (Supporting Information).

### SLM with oscillating carrying capacity

We considered the following stochastic differential equation as a model of the ecological dynamics of each community member

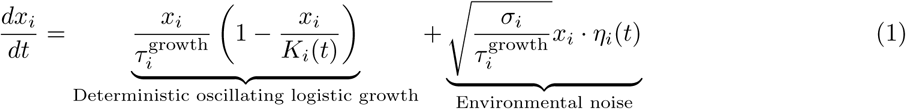

Here K*_i_*(t), σ*_i_*, and 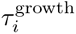 represent the carrying capacity, the strength of environmental noise, and the timescale of growth, respectively. This equation is known as the Stochastic Logistic Model of growth (SLM) [53], where we extended its original form by considering the carrying capacity as a function of time rather than as a constant. Motivated by the oscillatory nature of the logarithm of read counts and prior ecological modeling efforts [112, 113], we considered the following carrying capacity function.

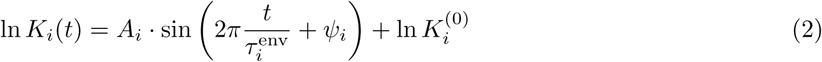

where the constants A*_i_*, 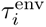, and ψ*_i_* specify the amplitude, inverse frequency, and phase of the sine function, respectively. The model reduces to a constant carrying capacity as 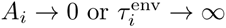, the latter of which can be interpreted as the timescale of environmental oscillations being far greater than our window of observation. The time-dependent solution of this form of the SLM is difficult to solve. Given that our oscillations occurred on a timescale far longer than the timescale of *∼* weekly sampling (i.e., 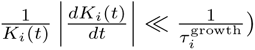, we modeled our data using the stationary probability distribution of the SLM as

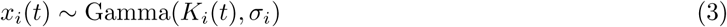

This distribution was used to derive the temporal autocorrelation of rRNA and rDNA as well as predictions of 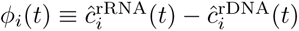 (Supporting Information).

For simulations, read counts were modeled as a multinomial sampling process

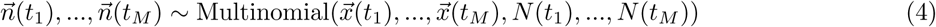

where 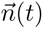 denotes the species abundance distribution for S species at time t*_m_*, M is the total number of samples, and N(t*_m_*) represents the total number of reads at time t*_m_*.

### Identifying an appropriate data transformation

Microbial 16S rRNA abundance data are typically examined as the relative abundance of read counts, defined as

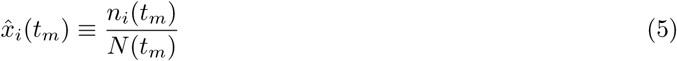

The presence of the total number of reads in the denominator results in this measure being subject to the effects of compositionality, where the constraint that the read counts of all community members at a given sample must sum to a fixed integer induces artifacts (e.g., oscillations) when there is none in the true abundances. Alternative measures that limit the effects of compositionality have been identified and effectively applied to 16S rRNA data (e.g., [114, 115]), one of which is the centered log-ratio (CLR). For this study we focused on the subset S*^∗^* of community members that had read counts greater than zero in all samples.

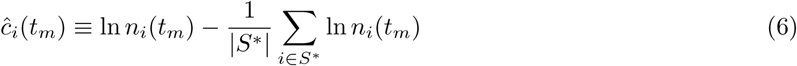

where *|* S*^∗^|* = 21 ASVs due to the high level of sequencing that was performed (i.e., sampling effort, Fig. S3). We validated this choice of transformation by comparing the accuracy of inferred sine parameters with a standard CLR-transformation using simulated read counts (Supporting Information).

### Statistical inference

#### Maximum likelihood inference of oscillatory parameters

We fit Eq. 2 using maximum likelihood to CLR-transformed data, constituting the ansatz 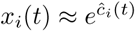. The mean time-dependent abundance and the squared inverse coefficient of variation can be obtained using the SLM parameters

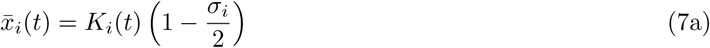

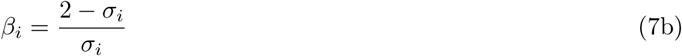

The distribution is then

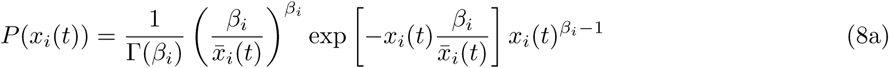

from which the log-likelihood function can be obtained

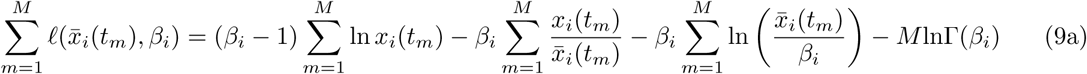

The parameters in x̄*_i_*(t) were inferred numerically using a two-step process. First, we performed a brute-force grid-search over ranges of parameter combinations and then used the parameter combination with the lowest negative log-likelihood as the initial parameters for optimization, using the likelihood as the objective function. Maximum likelihood inference was performed using lmfit [116]. Additional details are provided in the Supporting Information.

### A physiological model of rRNA:rDNA

To investigate the relationship between operon copy number and rRNA:rDNA and the rRNA-rDNA CPSD, we derived the following system of coupled SDEs:

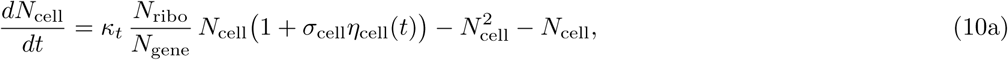

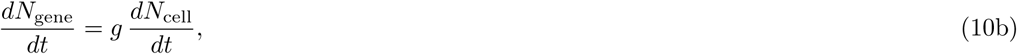

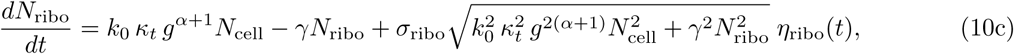

Here 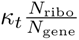 is the instantaneous *maximum* rate of per-capita growth, where the full per-capita growth function is analogous to the deterministic component of the SLM. The rate of per-capita ribosome assembly and degradation is k_0_ *·* g*^α^*^+1^ and γ, respectively, where α represents the gene dosage effect and k_0_ is a conversion rate.

The relationship between operon copy number g and steady-state rRNA:rDNA is then

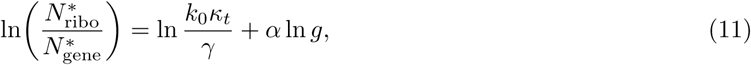

Mean rRNA operon copy numbers were obtained at the family level using the Ribosomal RNA Operon Copy Number Database (rrnDB) v5.8 [117].

By deriving the phase delay, we found that rRNA *precedes* rDNA when 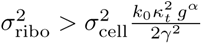. This result can be interpreted as the excess assembly of ribosomes relative to degradation being greater than the ribosomal-to-cellular noise ratio. Derivations are provided in the Supporting Information.

### Environmental variable analysis

We used Generalized Additive Models (GAMs) to model the relationship between our environmental variables and CLR-transformed abundances due to their flexibility and ability to handle non-linearity [118] using the R package mgcv [119]. We defined “sensitivity” as the derivative of the predicted CLR-transformed abundance with respect to a given environmental variable (Supporting Information).

## Data and code availability

All raw sequence data used in this study were obtained from a previous study [15]. Reprocessed data are available on Zenodo, DOI: 10.5281/zenodo.7692046. All code written for this study is available on GitHub under a GNU General Public License: rRNA_rDNA

## Acknowledgments

We thank A.M. Walczak for their insight on initial analyses and M. Cosentino Lagomarsino for their suggestions. We thank A. Goyal, J. Pasqualini, and N.I. Wisnoski for their comments on the manuscript. This work was supported by the NSF Postdoctoral Research Fellowships in Biology Program under Grant No. 2010885 (W.R.S.), Fondo Italiano per la Scienza - FIS (CUP J53C23002290001; J.G.).

## Author contributions

W.R.S., M.D.B., and J.G. conceptualized the project and wrote the manuscript. W.R.S. and J.G. performed all derivations. W.R.S. performed all analyses.

