## Supporting Information for "Leveraging rDNA and rRNA dynamics to investigate the link between ecology and physiology in a freshwater microbial community"

#### Supplemental materials and methods

##### Data acquisition

In the original study the epilimnion of a meso-eutrophic freshwater reservoir in Indiana, USA (39°11' N, 86°30' W) was sampled weekly from April 2013 to September 2015. Microbial biomass was filtered and V4 (primers 515F and 806R) 16S rRNA amplicon 250 x 250bp paired-end libraries were prepared and sequenced on an Illumina MiSeq using extracted DNA. To examine the community composition of rRNA, complementary DNA was obtained via Reverse-Transcription PCR, which was then sequenced using the same protocol as rDNA. The environmental variables temperature (°C), pH, specific conductivity (mS/cm), dissolved oxygen (mg/L), salinity (PSS), Secchi depth (m), total nitrogen (mg/L), total phosphorus ($\mu$g/L), and dissolved organic carbon (mg/L) were sampled at the same time as biomass. Additional information can be found in the original study (1). Throughout the manuscript we refer to the 16S rRNA amplicon sequence data for DNA and cDNA as rDNA and rRNA, respectively.

##### Data acquisition

We reprocessed raw FASTQ files in order to obtain Amplicon Sequence Variants (ASVs), the finest taxonomically resolved scale permitted by 16S rRNA sequencing. Raw FASTQ files were inspected for primers using `cutadapt v2.6` (2), where none were found. Correspondence with the authors of the original study revealed that primers were removed by the sequencing center. ASVs were then inferred using `dada2 v1.26` (3) in `R v4.5.1`. Raw paired-end reads were quality-filtered and trimmed using the `filterAndTrim()` funciton. Forward and reverse reads were truncated to 230 and 200 bp, respectively. Reads containing ambiguous bases (maxN = 0) or falling below quality thresholds (`maxEE = 2` and `maxEE = 5` for forward and reverse reads, respectively; `truncQ = 2`) were discarded, and PhiX reads were removed. Error rates were estimated separately for forward and reverse reads using `learnErrors()` and visually inspected via error profile plots. Denoising was performed using pooled sample inference to maximize detection of rare ASVs. Denoised forward and reverse reads were merged with `mergePairs()`, and per-sample sequence tables were constructed with `makeSequenceTable()`. Chimeric sequences were then identified and removed using `removeBimeraDenovo()` with the consensus method. Taxonomy was assigned using `v138.2` of Silva NR 99 for genus-level (`silva_nr99_v138.2_toGenus_trainset.fa.gz`) and species-level reference databases (`silva_v138.2_assignSpecies.fa.gz`). rRNA samples may have different error rates due to the data generation requiring an additional RT-PCR step to obtain cDNA. Therefore, ASVs from rRNA and rDNA samples were first inferred independently and then merged downstream. The fraction of reads that passed `dada2` inference and filtering was 91.2% and 83.9% for rDNA and rRNA, respectively.

##### Pairwise phylogenetic distances

Phylogenetic inference was performed on the subset of ASVs used in our analysis. ASVs were first aligned using `MUSCLE v5.3` with the `-super5` flag (4). Sites containing $\geq$ 80% uninformative bases were removed from the alignment. A phylogeny was inferred using `RAxML-NG v2.0.1` using a general time-reversible model with gamma-distributed variation in rates with the numbet of bootstraps determined using the `-autoMRE` flag (5). The 16S rRNA sequence from *Prochlorococcus marinus* subsp. *marinus* str. CCMP1375 was used as an outgroup (6). Pairwise phylogenetic distances were calculated using `ETE3 v3.1.2` (7).

#### Identifying an appropriate data transformation

The CLR-transformation as-is cannot handle zeros. One common solution is to add a pseudocount to all observations, though it is often unclear *a priori* whether this is a reasonable solution as it may introduce systematic biases to the data. We compared two forms of the CLR-transformation in order to identify a transformation that permits accurate inference of sine parameters. First, we used the entire set $S$ of ASVs ($|S| = 134,265$) by adding a pseudocount of $\epsilon = 1$ to all abundances.

$$\hat{c}_i^{(S)}(t_m) \equiv \ln\left(n_i(t_m) + \epsilon\right) - \frac{1}{S}\sum_{i=1}^{S} \ln\left(n_i(t_m) + \epsilon\right) \tag{S1}$$

Second, we identified the subset $S^*$ that had read counts greater than zero in all samples ($|S^*| = 25$) and performed the CLR-transformation on that subset.

$$\hat{c}_i(t_m) \equiv \ln\left(n_i(t_m)\right) - \frac{1}{|S^*|}\sum_{i \in S^*} \ln n_i(t_m) \tag{S2}$$

We investigated whether ASVs that were not sampled at every timepoint (i.e., occupancy $< 1$) had higher standard deviation of CLR-transformed abundance. Because the CLR-transformation is applied *per sample*, it is necessary to identify the subset of ASVs present in the same subset of samples so that the interpretation of the transformation is consistent across ASVs, effectively a double filtering process. We first identified the subset of samples where the same ASVs were consistently sampled. The CLR transform was then applied to each sample in this subset, allowing us to identify whether ASVs with lower occupancy across *all* samples displayed higher variation. The coefficient of variation is not an appropriate measure of temporal variation in this instance because CLR-transformed timeseries regularly fluctuate above and below zero, meaning that the absolute value of the mean can be arbitrarily close to zero, blowing up the CV. Instead, we normalized the standard deviation of $\hat{c}_i$ by the *mean of the absolute value*, $\overline{|\hat{c}_i|}$.

#### Maximum likelihood inference of ASVs

Given our focus on ASVs that were detected in all samples, we fit Eq. 4 to the CLR-transformed data, constituting the ansatz $x_i(t) \propto e^{\hat{c}_i(t)}$. The variable $x_i$ follows the Stochastic Logistic Model (SLM) with time-varying carrying capacity

$$\frac{dx_i}{dt} = \frac{x_i}{\tau_i^{\text{growth}}}\left(1 - \frac{x_i}{K_i(t)}\right) + \sqrt{\frac{\sigma_i}{\tau_i^{\text{growth}}}}x_i \cdot \eta_i(t) \tag{S3}$$

When the the fractional rate of change of the carrying capacity is slow relative to the rate of growth, $\frac{1}{K_i(t)}\left|\frac{dK_i(t)}{dt}\right| \ll \frac{1}{\tau_i^{\text{growth}}}$, then we can invoke the quasi-stationary approximation to the Fokker-Planck form of the SLM, obtaining a gamma distribution with time-varying mean.

$$P(x_i) = \frac{1}{\Gamma(\beta_i)}\left(\frac{\beta_i}{\bar{x}_i}\right)^{\beta_i} \exp\left[-x_i\frac{\beta_i}{\bar{x}_i}\right] x_i^{\beta_i - 1} \tag{S4a}$$

where

$$\bar{x}_i = K_i\left(1 - \frac{\sigma_i}{2}\right) \tag{S5a}$$

$$\beta_i = \frac{2 - \sigma_i}{\sigma_i} \tag{S5b}$$

We then numerically maximized the likelihood to infer the parameters of our oscillating gamma distribution. If the mean is a constant, the above distribution has the following log-likelihood function

$$\ell(\bar{x}_i, \beta_i) = \sum \ln P(x_i|\bar{x}_i, \beta_i) \tag{S6a}$$

$$= (\beta_i - 1)\sum_{m=1}^{M} \ln x_i(t_m) - \frac{\beta_i}{\bar{x}_i}\sum_{m=1}^{M} x_i(t_m) - M\beta_i \ln\left(\frac{\bar{x}_i}{\beta_i}\right) - M\ln\Gamma(\beta_i) \tag{S6b}$$

By taking the partial derivative of $\ell$ with respect to $\beta$ we find that the maximum likelihood solution for $\beta_i$ is the solution to the following equation

$$\ln(\beta_i) - \psi_0(\beta_i) = \ln\left(\frac{1}{M}\sum_{m=1}^{M} x_i(t_m)\right) - \frac{1}{M}\sum_{m=1}^{M} \ln x_i(t_m) \tag{S7}$$

where $\psi_0(\cdot)$ is the digamma function. The expression can be solved numerically, from which we calculate $\sigma_i$.

For our oscillating mean model the likelihood function has the following form, where the mean is now a time-dependent parameter

$$\sum_{m=1}^{M} \ell(\bar{x}_i(t_m), \beta_i) = (\beta_i - 1)\sum_{m=1}^{M} \ln x_i(t_m) - \beta_i \sum_{m=1}^{M} \frac{x_i(t_m)}{\bar{x}_i(t_m)} - \beta_i \sum_{m=1}^{M} \ln\left(\frac{\bar{x}_i(t_m)}{\beta_i}\right) - M\ln\Gamma(\beta_i) \tag{S8a}$$

The partial derivatives with respect to the mean and $\beta$ is

$$\sum_{m=1}^{M} \frac{\partial \ell(\bar{x}_i(t_m), \beta_i)}{\partial \bar{x}_i(t_m)} = \beta_i \sum_{m=1}^{M} \frac{x_i(t_m)}{\bar{x}_i(t_m)^2} - \beta_i \sum_{m=1}^{M} \frac{1}{\bar{x}_i(t_m)} \tag{S9a}$$

$$\sum_{m=1}^{M} \frac{\partial \ell(\bar{x}_i(t_m), \beta_i)}{\partial \beta} = \sum_{m=1}^{M} \ln x_i(t_m) - \sum_{m=1}^{M} \frac{x_i(t_m)}{\bar{x}_i(t_m)} - \sum_{m=1}^{M} \ln \bar{x}_i(t_m) + M(\ln(\beta_i) + 1) - M\psi_0(\beta_i) \tag{S9b}$$

For the parameters of the oscillating carrying capacity, we find the partial derivatives of the mean with respect to each parameter

$$\frac{\partial \bar{x}_i(t)}{\partial K_0} = \exp\left[A \cdot \sin\left(\frac{2\pi}{\tau_{\text{env}}}t + \psi\right)\right] \tag{S10a}$$

$$\frac{\partial \bar{x}_i(t)}{\partial \tau_{\text{env}}} = -\frac{K_0}{\tau_{\text{env}}^2} A 2\pi t \cdot \exp\left[A \cdot \sin\left(\frac{2\pi}{\tau_{\text{env}}}t + \psi\right)\right] \cdot \cos\left(\frac{2\pi}{\tau_{\text{env}}}t + \psi\right) \tag{S10b}$$

$$\frac{\partial \bar{x}_i(t)}{\partial \psi} = K_0 A \cdot \exp\left[A \cdot \sin\left(\frac{2\pi}{\tau_{\text{env}}}t + \psi\right)\right] \cdot \cos\left(\frac{2\pi}{\tau_{\text{env}}}t + \psi\right) \tag{S10c}$$

$$\frac{\partial \bar{x}_i(t)}{\partial A} = K_0 \cdot \exp\left[A \cdot \sin\left(\frac{2\pi}{\tau_{\text{env}}}t + \psi\right)\right] \cdot \sin\left(\frac{2\pi}{\tau_{\text{env}}}t + \psi\right) \tag{S10d}$$

The MLE solution for each parameter in the mean is

$$\sum_{m=1}^{M} \frac{\partial \ell(\bar{x}_i(t_m),\beta_i)}{\partial \bar{x}_i(t_m)} \frac{\partial \bar{x}_i(t_m)}{\partial K_0} = \beta_i \sum_{m=1}^{M} \frac{x_i(t_m)}{\bar{x}_i(t_m)^2} \frac{\partial \bar{x}_i(t_m)}{\partial K_0} - \beta_i \sum_{m=1}^{M} \frac{1}{\bar{x}_i(t_m)} \frac{\partial \bar{x}_i(t_m)}{\partial K_0} = 0 \tag{S11a}$$

$$\sum_{m=1}^{M} \frac{\partial \ell(\bar{x}_i(t_m),\beta_i)}{\partial \bar{x}_i(t_m)} \frac{\partial \bar{x}_i(t_m)}{\partial \tau_{\mathrm{env}}} = \beta_i \sum_{m=1}^{M} \frac{x_i(t_m)}{\bar{x}_i(t_m)^2} \frac{\partial \bar{x}_i(t_m)}{\partial \tau_{\mathrm{env}}} - \beta_i \sum_{m=1}^{M} \frac{1}{\bar{x}_i(t_m)} \frac{\partial \bar{x}_i(t_m)}{\partial \tau_{\mathrm{env}}} = 0 \tag{S11b}$$

$$\sum_{m=1}^{M} \frac{\partial \ell(\bar{x}_i(t_m),\beta_i)}{\partial \bar{x}_i(t_m)} \frac{\partial \bar{x}_i(t_m)}{\partial \psi} = \beta_i \sum_{m=1}^{M} \frac{x_i(t_m)}{\bar{x}_i(t_m)^2} \frac{\partial \bar{x}_i(t_m)}{\partial \psi} - \beta_i \sum_{m=1}^{M} \frac{1}{\bar{x}_i(t_m)} \frac{\partial \bar{x}_i(t_m)}{\partial \psi} = 0 \tag{S11c}$$

$$\sum_{m=1}^{M} \frac{\partial \ell(\bar{x}_i(t_m),\beta_i)}{\partial \bar{x}_i(t_m)} \frac{\partial \bar{x}_i(t_m)}{\partial A} = \beta_i \sum_{m=1}^{M} \frac{x_i(t_m)}{\bar{x}_i(t_m)^2} \frac{\partial \bar{x}_i(t_m)}{\partial A} - \beta_i \sum_{m=1}^{M} \frac{1}{\bar{x}_i(t_m)} \frac{\partial \bar{x}_i(t_m)}{\partial A} = 0 \tag{S11d}$$

From which we obtain

$$\sum_{m=1}^{M} x_i(t_m) \cdot \exp\left[A \cdot \sin\left(\frac{2\pi}{\tau_{\mathrm{env}}} t + \psi\right)\right]^{-1} - M K_0 = 0 \tag{S12a}$$

$$\sum_{m=1}^{M} t_m \cdot \frac{x_i(t_m)}{\bar{x}_i(t_m)} \cos\left(\frac{2\pi}{\tau_{\mathrm{env}}} t_m + \psi\right) - \sum_{m=1}^{M} t_m \cdot \cos\left(\frac{2\pi}{\tau_{\mathrm{env}}} t_m + \psi\right) = 0 \tag{S12b}$$

$$\sum_{m=1}^{M} \frac{x_i(t_m)}{\bar{x}_i(t_m)} \cos\left(\frac{2\pi}{\tau_{\mathrm{env}}} t_m + \psi\right) - \sum_{m=1}^{M} \cos\left(\frac{2\pi}{\tau_{\mathrm{env}}} t_m + \psi\right) = 0 \tag{S12c}$$

$$\sum_{m=1}^{M} \frac{x_i(t_m)}{\bar{x}_i(t_m)} \sin\left(\frac{2\pi}{\tau_{\mathrm{env}}} t_m + \psi\right) - \sum_{m=1}^{M} \sin\left(\frac{2\pi}{\tau_{\mathrm{env}}} t_m + \psi\right) = 0 \tag{S12d}$$

We can rearrange the first equation to obtain $K_0$ as a function of the remaining parameters, reducing the system of equations to three. The derivative of the log-likelihood function would then have to be solved simultaneously for all parameters in $\bar{x}_i$. There is no analytic solution, so the parameters in $\bar{x}_i$ as well as $\beta_i$ were inferred numerically by minimizing the negative log-likelihood using the `Scipy` function minimize with the L-BFGS-B algorithm to place bounds on the parameters. However, our numerical approach lead to overfitting of $\bar{x}_i(t)$ that resulted in $\beta_i \to 0$, corresponding to $\sigma_i \to \infty$, a limit that cannot occur for the SLM since as it requires $\sigma_i < 2$

As a solution, we elected to perform a two-step fitting procedure. First, we made the assumption that the relation in Eq. S7 approximately holds when the mean is oscillating, the validity of which we tested by performing simulations with varying oscillatory parameters and $\sigma_i$ (Fig. **??**). Second, we identified parameters that minimized the negative log-likelihood by 1) performing a brute-force grid-search over ranges of parameter combinations and then 2) using the parameter combination with the lowest negative log-likelihood as the initial parameters for optimization with the likelihood as the objective function. Maximum likelihood inference was performed using `lmfit` (8).

We then updated our initial inferred value of $\beta_i$ using the inferred parameters of the time-varying mean. The rationale of this decision was that the maximum likelihood estimator of $\beta_i$ (Eq. S7) receives a positive contribution from the temporal variation in $\bar{x}_i(t)$ that is independent of the true $\beta_i$. Specifically, since $\bar{x}_i(t) \propto e^{A_i \sin(\cdot)}$, the time-average of $\bar{x}_i(t)$ exceeds the geometric mean. This excess inflates the right-hand side of Eq. S7, implying a smaller solution $\hat{\beta}_i$ and therefore a larger $\hat{\sigma}_i$, causing the noise to be overestimated, which alters the predicted temporal autocorrelation of $\ln x_i(t)$.

To obtain an estimate of $\beta_i$ free from this bias, we exploited the fact that the variance of $\ln x_i(t)$ about its conditional expectation $\langle \ln x_i(t) \rangle$ depends only on $\beta_i$ and *not* on the time-varying mean. That is, for our gamma distribution, $\mathrm{Var}[\ln x_i(t) - \ln \bar{x}_i(t)] = \psi_1(\beta_i)$. After obtaining maximum likelihood estimates of the mean parameters, we computed the log-residuals

$$r_m \ = \ \ln x_i(t_m) - \ln \bar{x}_i(t_m), \qquad m = 1, \ldots, M, \tag{S13}$$

where $\bar{x}_i(t_m)$ is the fitted mean evaluated at time $t_m$. The residuals $r_m$ have mean $\psi_0(\beta_i) - \ln \beta_i$ and variance $\psi_1(\beta_i)$. We therefore obtained a revised estimate $\hat{\beta}_i^{\mathrm{corr}}$ by numerically solving:

$$\psi_1(\beta_i^{\mathrm{corr}}) = \frac{1}{M} \sum_{m=1}^{M} (r_m - \bar{r})^2 \,, \tag{S14}$$

using Brent's root-finding method. Because $\psi_1$ is strictly positive and monotonically decreasing on $(0, \infty)$, Eq. S14 has a unique solution provided $\mathrm{Var}[r_m] > 0$. The revised noise parameter estimate was used in place of the original two-step estimate for all downstream analyses.

We assessed significance of our fit for each ASV via parametric bootstrapping. Specifically, we repeated our numerical maximum likelihood procedure for a gamma with a constant mean and compared its likelihood to that of our model of a gamma with time-varying mean. For each ASV we simulated $10^4$ timeseries using our inferred parameters from a gamma with constant mean. For each timeseries we fit both the time-varying and constant mean gamma models and computed the log-likelihood ratio, obtaining a null distribution of likelihood ratios. We then calculated $P$-values using this null distribution as a reference.

##### Ruling out potential confounders in the data

Because aquatic communities are often diurnal, we confirmed that variation in intra-daily sampling times was unlikely to shape our oscillation inference as there was no correlation between sampling time and the day of the year on which a sample was collected (Fig. S7). Because we subset the community to ASVs present in all samples, we tested whether rarer ASVs displayed greater temporal variation in CLR-transformed abundance (i.e., boom-bust dynamics) and found no evidence (Fig. S8).

We calculated $\Delta\psi_i \equiv (\psi_i^{\mathrm{rRNA}} - \psi_i^{\mathrm{rDNA}} + \pi) \mathrm{mod}(2\pi) - \pi$ so that $\Delta\psi_i$ has the domain $[-\pi, \pi)$. We tested for non-uniformity in $\Delta\psi$ using the Rayleigh test on the unit circle (9). The mean resultant vector plotted in Fig. SS16 was calculated as

$$\mathbf{R} = \frac{1}{N} \sum_{i=1}^{N} \begin{pmatrix} \cos \Delta\psi_i \\ \sin \Delta\psi_i \end{pmatrix}$$

where the statistic $R$ is the magnitude of the vector

$$R = \|\mathbf{R}\|,$$

##### Least-squares inference of environmental variables

We view the dynamics of measured environmental variables as effectively deterministic by fitting sinusoidal functions to the data rather than with a probability distribution. We used the same procedure as our MLE inference of ASV dynamics, but with the least-squares difference as the objective function.

##### Autocorrelation of the gamma with time-varying mean for rRNA and rDNA

The autocorrelation of a gamma random variable with time-varying mean can be derived for rRNA and rDNA. Because we are interpreting the CLR-transformation as a log-transform that accounts for compositionality, we must derive the autocorrelation of the log of the random variable. To simplify the derivation, we decompose the mean into time-dependent and independent components.

$$\big\langle \ln x_i(t) \big\rangle = J_i + \tilde{K}_i(t), \qquad J_i \;\equiv\; \psi_0(\beta_i) - \ln\beta_i + \ln\!\big(1 - \tfrac{\sigma_i}{2}\big) + \ln K_i^{(0)}, \qquad \tilde{K}_i(t) \equiv A_i \cdot \sin\left(2\pi \frac{t}{\tau_i^{\mathrm{env}}} + \psi_i\right). \tag{S15}$$

where $\psi_0(\cdot)$ is the digamma function. The stochastic fluctuation about this mean is:

$$\varepsilon_i(t) \;\equiv\; \ln x_i(t) - \big\langle \ln x_i(t) \big\rangle, \qquad \big\langle \varepsilon_i(t) \big\rangle = 0, \qquad \big\langle \varepsilon_i(t)^2 \big\rangle = \psi_1(\beta_i) \tag{S16}$$

where $\psi_1(\cdot)$ is the trigamma function. The phase $\psi_i \sim \mathrm{Uniform}[-\pi, \pi)$ is treated as random to confer stationarity. Writing $\ln x_i(t) = J_i + \tilde{K}_i(t) + \varepsilon_i(t)$, the autocovariance at lag $\delta t = t_2 - t_1$ is:

$$R_{\ln x_i}(\delta t) = \Big\langle \big(\ln \bar{x}_i(t_1) - J_i\big)\big(\ln \bar{x}_i(t_2) - J_i\big) \Big\rangle \tag{S17a}$$

$$= \Big\langle \big[\tilde{K}_i(t_1) + \varepsilon_i(t_1)\big]\,\big[\tilde{K}_i(t_2) + \varepsilon_i(t_2)\big] \Big\rangle \tag{S17b}$$

$$= \big\langle \tilde{K}_i(t_1)\,\tilde{K}_i(t_2) \big\rangle + \big\langle \tilde{K}_i(t_1)\,\varepsilon_i(t_2) \big\rangle + \big\langle \varepsilon_i(t_1)\,\tilde{K}_i(t_2) \big\rangle + \big\langle \varepsilon_i(t_1)\,\varepsilon_i(t_2) \big\rangle. \tag{S17c}$$

Since $\varepsilon_i(t)$ is zero-mean and independent of the deterministic component $\tilde{K}_i(t)$, the cross terms vanish, giving us:

$$R_{\ln x_i}(\delta t) = \big\langle \tilde{K}_i(t_1)\,\tilde{K}_i(t_2) \big\rangle + \big\langle \varepsilon_i(t_1)\,\varepsilon_i(t_2) \big\rangle \tag{S18}$$

Solving the first term, we obtain

$$\big\langle \tilde{K}_i(t_1)\,\tilde{K}_i(t_2) \big\rangle = \big\langle A_i \sin\!\Big(2\pi \tfrac{t_1}{\tau_i^{\mathrm{env}}} + \psi_i\Big)\, A_i \sin\!\Big(2\pi \tfrac{t_2}{\tau_i^{\mathrm{env}}} + \psi_i\Big) \big\rangle \tag{S19a}$$

$$= \frac{A_i^2}{2\pi} \int_{-\pi}^{\pi} \sin\!\left(2\pi \frac{t_1}{\tau_i^{\mathrm{env}}} + \psi_i\right) \sin\!\left(2\pi \frac{t_2}{\tau_i^{\mathrm{env}}} + \psi_i\right) d\psi_i \tag{S19b}$$

$$= \frac{A_i^2}{4\pi} \int_{-\pi}^{\pi} \left[\cos\!\left(2\pi \frac{t_1 - t_2}{\tau_i^{\mathrm{env}}}\right) - \cos\!\left(2\pi \frac{t_1 + t_2}{\tau_i^{\mathrm{env}}} + 2\psi_i\right)\right] d\psi_i \tag{S19c}$$

$$= \frac{A_i^2}{2} \cos\!\left(2\pi \frac{t_1 - t_2}{\tau_i^{\mathrm{env}}}\right), \tag{S19d}$$

Where we have used the identity $\sin\alpha \sin\beta = \frac{1}{2}[\cos(\alpha - \beta) - \cos(\alpha + \beta)]$. Assuming $\varepsilon_i(t)$ is uncorrelated across time:

$$\big\langle \varepsilon_i(t_1)\,\varepsilon_i(t_2) \big\rangle = \psi_1(\beta_i)\ \cdot \mathbf{1}[t_1 = t_2], \tag{S20}$$

We then normalize by $R_{\ln x_i}(0)$ to obtain the autocorrelation

$$\rho_{\ln x_i}(\delta t) = \frac{R_{\ln x_i}(\delta t)}{R_{\ln x_i}(0)} \tag{S21a}$$

$$= \frac{\dfrac{A_i^2}{2} \cos\!\left(2\pi \dfrac{\delta t}{\tau_i^{\mathrm{env}}}\right) + \psi_1(\beta_i)\ \mathbf{1}[\delta t = 0]}{\dfrac{A_i^2}{2} + \psi_1(\beta_i)}. \tag{S21b}$$

We compared predicted autocorrelations to those calculated from CLR-transformed timeseries (Figs. S11, S12).

#### Autocorrelation of the gamma with time-varying mean for rRNA:rDNA

We calculated the difference between CLR-transformed rRNA and rDNA abundances as a proxy for the rRNA:rDNA ratio. The expected CLR-transformed abundances of ASVs follow the sine function, so we can predict the expected difference $\left\langle \hat{c}_i^{\text{rRNA}}(t) - \hat{c}_i^{\text{rDNA}}(t) \right\rangle$ as the difference between two oscillating carrying capacities, known as the phase difference. Using our model of a gamma distribution with time-varying mean, we obtain the following expression

$$\phi_i(t) \;\equiv\; \ln x_i^{\text{rRNA}}(t) - \ln x_i^{\text{rDNA}}(t) \;=\; \Delta J_i + \underbrace{\tilde{K}_i^{\text{rRNA}}(t) - \tilde{K}_i^{\text{rDNA}}(t)}_{\Delta \tilde{K}_i(t)} + \underbrace{\varepsilon_i^{\text{rRNA}}(t) - \varepsilon_i^{\text{rDNA}}(t)}_{\xi_i(t)}, \tag{S22}$$

where $\Delta J_i = J_i^{\text{rRNA}} - J_i^{\text{rDNA}}$ is a constant and the combined noise $\xi_i(t)$ satisfies

$$\langle \xi_i(t) \rangle = 0, \qquad \langle \xi_i(t_1)\, \xi_i(t_2) \rangle = \underbrace{\left[\psi_1(\beta_i^{\text{rRNA}}) + \psi_1(\beta_i^{\text{rDNA}})\right]}_{\Psi_i} \cdot \mathbf{1}[t_1 = t_2]. \tag{S23}$$

The autocovariance is then

$$\begin{aligned} R_{\phi_i}(\delta t) &= \Big\langle \big(\phi_i(t_1) - \Delta J_i\big)\big(\phi_i(t_2) - \Delta J_i\big) \Big\rangle && \text{(S24a)} \\ &= \Big\langle \big[\Delta \tilde{K}_i(t_1) + \xi_i(t_1)\big]\big[\Delta \tilde{K}_i(t_2) + \xi_i(t_2)\big] \Big\rangle && \text{(S24b)} \\ &= \langle \Delta \tilde{K}_i(t_1)\, \Delta \tilde{K}_i(t_2) \rangle + \langle \xi_i(t_1)\, \xi_i(t_2) \rangle, && \text{(S24c)} \end{aligned}$$

where cross terms $\langle \Delta \tilde{K}_i(t)\, \xi_i(t') \rangle = 0$ vanish. We expand the first product to obtain

$$\begin{aligned} \langle \Delta \tilde{K}_i(t_1)\, \Delta \tilde{K}_i(t_2) \rangle = {} & \langle \tilde{K}_i^{\text{rRNA}}(t_1)\, \tilde{K}_i^{\text{rRNA}}(t_2) \rangle - \langle \tilde{K}_i^{\text{rRNA}}(t_1)\, \tilde{K}_i^{\text{rDNA}}(t_2) \rangle \\ & - \langle \tilde{K}_i^{\text{rDNA}}(t_1)\, \tilde{K}_i^{\text{rRNA}}(t_2) \rangle + \langle \tilde{K}_i^{\text{rDNA}}(t_1)\, \tilde{K}_i^{\text{rDNA}}(t_2) \rangle. \end{aligned} \tag{S25}$$

Cross terms vanish as $\psi_i^{\text{rRNA}}$ and $\psi_i^{\text{rDNA}}$ are independent, so the phase-averaged sines are zero

$$\langle \tilde{K}_i^{\text{rRNA}}(t_1)\, \tilde{K}_i^{\text{rDNA}}(t_2) \rangle = A_i^{\text{rRNA}} A_i^{\text{rDNA}} \underbrace{\left\langle \sin\!\left(\tfrac{2\pi t_1}{\tau_i^{\text{rRNA}}} + \psi_i^{\text{rRNA}}\right) \right\rangle_{\psi^{\text{rRNA}}}}_{=0} \cdot \left\langle \sin\!\left(\tfrac{2\pi t_2}{\tau_i^{\text{rDNA}}} + \psi_i^{\text{rDNA}}\right) \right\rangle_{\psi^{\text{rDNA}}} = 0. \tag{S26}$$

Therefore, only the auto-terms remain

$$\begin{aligned} \langle \tilde{K}_i^{\text{rRNA}}(t_1)\, \tilde{K}_i^{\text{rRNA}}(t_2) \rangle &= \frac{(A_i^{\text{rRNA}})^2}{2} \cos\!\left(\frac{2\pi\, \delta t}{\tau_i^{\text{rRNA}}}\right) && \text{(S27a)} \\ \langle \tilde{K}_i^{\text{rDNA}}(t_1)\, \tilde{K}_i^{\text{rDNA}}(t_2) \rangle &= \frac{(A_i^{\text{rDNA}})^2}{2} \cos\!\left(\frac{2\pi\, \delta t}{\tau_i^{\text{rDNA}}}\right) && \text{(S27b)} \end{aligned}$$

Combining all terms with $\delta t = t_1 - t_2$, we obtain the final autocovariance expression

$$R_{\phi_i}(\delta t) = \frac{(A_i^{\mathrm{rRNA}})^2}{2}\cos\left(\frac{2\pi\,\delta t}{\tau_i^{\mathrm{rRNA}}}\right) + \frac{(A_i^{\mathrm{rDNA}})^2}{2}\cos\left(\frac{2\pi\,\delta t}{\tau_i^{\mathrm{rDNA}}}\right) + \Psi_i\,\mathbf{1}[\delta t = 0] \tag{S28}$$

where $\Psi_i = \psi_1(\beta_i^{\mathrm{rRNA}}) + \psi_1(\beta_i^{\mathrm{rDNA}})$. From this result we can obtain the autocorrelation

$$\rho_{\phi_i}(\delta t) = \frac{\frac{(A_i^{\mathrm{rRNA}})^2}{2}\cos\left(\frac{2\pi\,\delta t}{\tau_i^{\mathrm{rRNA}}}\right) + \frac{(A_i^{\mathrm{rDNA}})^2}{2}\cos\left(\frac{2\pi\,\delta t}{\tau_i^{\mathrm{rDNA}}}\right)}{\frac{(A_i^{\mathrm{rRNA}})^2 + (A_i^{\mathrm{rDNA}})^2}{2} + \Psi_i} \tag{S29}$$

We note that this expression can be reduced to that of a single cosine term if rRNA and rDNA have identical oscillation timescales, $\tau_i^{\mathrm{rRNA}} = \tau_i^{\mathrm{rDNA}} = \tau_i$.

#### Simulating communities with oscillating ASVs

We simulated the scenario where an oscillating ASV impacted the relative abundances of non-oscillating ASVs. Specifically, we determined whether parameters in Eq. 4 could be accurately inferred for relative abundances along with CLR-transformed read counts where 1) a pseudocount of one was added to all ASVs in all samples or 2) only the subset of ASVs present in all samples was selected. Given that the mean relative abundance is proportional to the carrying capacity under the SLM and empirical mean relative abundances follow a lognormal distribution (10), we drew $K_i^{(0)}$ as lognormal random variables for $10^3$ ASVs. We assigned $\max[K_i^{(0)}]$ to the oscillating ASV. The $\sigma_i$ of each ASV was simulated as an exponentially distributed random variable based on previous results (11), where we manipulated the rate parameter. Relative abundances were independently sampled as gamma random variables using the same number of samples as in the empirical data as well as the time range. Read counts were sampled as a multinomial process using $10^5$ reads, a value similar to what was found in the dataset (Fig. S3). The two forms of the CLR were applied to the read counts along with the relative abundance. The model was fit using the same procedure as the empirical data to the one oscillating ASV which also had the highest rank abundance and to the second highest rank ASV which was non-oscillating.

##### The probability distribution of rRNA:rDNA

Barcode-based studies in microbial ecology and environmental microbiology consistently interpret the ratio between rRNA and rDNA relative abundances (i.e., rRNA:rDNA) as a proxy for metabolic activity (12; 13; 14; 15; 16; 17; 18; 19). The underlying assumption here is that rRNA:rDNA is reflective of ribosomal content. There is also an additional, statistical assumption: that the ratio contains information that is otherwise absent from rRNA and rDNA alone.

We can predict the distribution of rRNA:rDNA values by recognizing that the AFDs of rDNA and rRNA, separately, are well-described by a gamma distribution. The full time-varying mean containing four parameters is unnecessary in this instance, as a constant mean was sufficient to fit a gamma to the empirical distributions. Given that we are treating $\hat{c}_i$ as a compositionality-corrected log relative abundance, we can predict rRNA:rDNA as the difference of two log-gamma distributed random variables.

$$X \equiv \hat{c}_i^{\mathrm{rDNA}} \sim \mathrm{Gamma}\left(\beta_i^{\mathrm{rDNA}},\, \frac{\bar{x}_i^{\mathrm{rDNA}}}{\beta_i^{\mathrm{rDNA}}}\right), \qquad Y \equiv \hat{c}_i^{\mathrm{rRNA}} \sim \mathrm{Gamma}\left(\beta_i^{\mathrm{rRNA}},\, \frac{\bar{x}_i^{\mathrm{rRNA}}}{\beta_i^{\mathrm{rRNA}}}\right),$$

Then rRNA:rDNA is defined as

$$Z \;\equiv\; \frac{Y}{X} \;=\; e^{\,\hat{c}_i^{\mathrm{rRNA}} - \hat{c}_i^{\mathrm{rDNA}}} \;=\; e^{\,\hat{\phi}_i}$$

To simplify notation we will define $\beta_X \equiv \beta_i^{\mathrm{rDNA}}$, $\beta_Y \equiv \beta_i^{\mathrm{rRNA}}$, $r_X \equiv \frac{\alpha_1}{\bar{x}_i^{\mathrm{rDNA}}}$, and $r_Y \equiv \frac{\alpha_2}{\bar{x}_i^{\mathrm{rRNA}}}$. We derive our distribution of interest by first marginalizing over $X$:

$$P_Z(z) = \int_0^\infty x \cdot P_Y(xz)\, P_X(x)\, \mathrm{d}x.$$

Substituting the two gamma distributions gives

$$\begin{aligned} P_Z(z) &= \int_0^\infty x \cdot \frac{r_Y^{\beta_Y}}{\Gamma(\beta_Y)} (xz)^{\beta_Y - 1} e^{-r_Y xz} \cdot \frac{r_X^{\beta_X}}{\Gamma(\beta_X)} x^{\beta_X - 1} e^{-r_X x}\, \mathrm{d}x \\ &= \frac{r_X^{\beta_X} r_Y^{\beta_Y}}{\Gamma(\beta_X)\Gamma(\beta_Y)}\, z^{\beta_Y - 1} \int_0^\infty x^{\alpha_X + \beta_Y - 1}\, e^{-(r_X + r_Y z)x}\, \mathrm{d}x \end{aligned}$$

We recognize that our remaining integral is a standard gamma function,

$$\int_0^\infty x^{\beta_X + \beta_Y - 1}\, e^{-(r_X + r_Y z)x}\, \mathrm{d}x = \frac{\Gamma(\beta_X + \beta_Y)}{(r_X + r_Y z)^{\beta_X + \beta_Y}}.$$

Using the definition of the beta function and factoring the denominator, we obtain:

$$P_Z(z) = \frac{1}{B(\beta_X, \beta_Y)} \cdot \frac{(r_Y/r_X)^{\beta_Y}\, z^{\beta_Y - 1}}{\big(1 + (r_Y/r_X)\, z\big)^{\beta_X + \beta_Y}}, \qquad z > 0. \tag{S30}$$

Which is the *Beta-prime distribution* scaled by $\lambda \equiv r_Y/r_X$ (20). We can now obtain the distribution of the *logarithm* of the ratio. Given that $\hat{\phi}_i = \log Z$, i.e. $Z = e^{\hat{\phi}_i}$, the change-of-variables Jacobian is $|\mathrm{d}z/\mathrm{d}\hat{\phi}_i| = e^{\hat{\phi}_i}$, giving

$$P(\hat{\phi}_i) = P_Z\left(e^{\hat{\phi}_i}\right) \cdot e^{\hat{\phi}_i} = \frac{1}{B(\beta_Y, \beta_X)} \cdot \frac{\lambda^{\beta_Y} \left(e^{\hat{\phi}_i}\right)^{\beta_Y - 1}}{\left(1 + \lambda\, e^{\hat{\phi}_i}\right)^{\beta_X + \beta_Y}} \cdot e^{\hat{\phi}_i}$$

Collecting the exponentials in the numerator, we obtain:

$$P(\hat{\phi}_i) = \frac{1}{B(\beta_Y, \beta_X)} \cdot \frac{\left(\lambda\, e^{\hat{\phi}_i}\right)^{\beta_Y}}{\left(1 + \lambda\, e^{\hat{\phi}_i}\right)^{\beta_X + \beta_Y}} \tag{S31}$$

Re-substituting our original variables and using the symmetry property of the beta function, we obtain:

$$P\left(\hat{\phi}_i\right) = \frac{1}{B\left(\beta_i^{\mathrm{rRNA}}, \beta_i^{\mathrm{rDNA}}\right)} \cdot \frac{\left(e^{\hat{\phi}_i} \cdot \dfrac{\bar{x}_i^{\mathrm{rDNA}}}{\bar{x}_i^{\mathrm{rRNA}}} \cdot \dfrac{\beta_i^{\mathrm{rRNA}}}{\beta_i^{\mathrm{rDNA}}}\right)^{\beta_i^{\mathrm{rRNA}}}}{\left(1 + e^{\hat{\phi}_i} \cdot \dfrac{\bar{x}_i^{\mathrm{rDNA}}}{\bar{x}_i^{\mathrm{rRNA}}} \cdot \dfrac{\beta_i^{\mathrm{rRNA}}}{\beta_i^{\mathrm{rDNA}}}\right)^{\beta_i^{\mathrm{rRNA}} + \beta_i^{\mathrm{rDNA}}}} \tag{S32}$$

This predicted distribution of $\hat{\phi}_i$ requires no fitting on the empirical distribution of $\hat{\phi}_i$, solely the empirical distributions of rRNA and rDNA.

##### Temporal cross-correlations of rRNA:rDNA

Given that rRNA:rDNA is often interpreted as being reflective of the ribosomal activity within a taxon, a key physiological parameter of growth, it is worth asking how $\hat{\phi}_i(t)$ relates to the change in $\hat{c}(t)_i^{\mathrm{rDNA}}$ at a subsequent timepoint, $\delta\hat{c}(t)_i^{\mathrm{rDNA}} \equiv \hat{c}(t+\delta t)_i^{\mathrm{rDNA}} - \hat{c}(t)_i^{\mathrm{rDNA}}$. However, estimated correlations between these two quantities are in danger of being spurious due to both variables sharing $\hat{c}(t)_i^{\mathrm{rDNA}}$. To illustrate this, consider the scenario where both timeseries are permuted, removing any underlying correlation between rRNA and rDNA. The estimated correlation between $\hat{\phi}_i(t)$ and $\delta\hat{c}(t)_i^{\mathrm{rDNA}}$ is then

$$\begin{aligned}
\mathrm{Corr}(\hat{\phi}_i(t), \delta\hat{c}(t)_i^{\mathrm{rDNA}}) &= \frac{\mathrm{Cov}(\hat{\phi}_i(t), \delta\hat{c}(t)_i^{\mathrm{rDNA}})}{\sqrt{\mathrm{Var}(\hat{\phi}_i(t)) \cdot \mathrm{Var}(\delta\hat{c}(t)_i^{\mathrm{rDNA}})}} && \text{(S33a)} \\
&= \frac{\mathrm{Cov}(-\hat{c}(t)_i^{\mathrm{rDNA}}, -\hat{c}(t)_i^{\mathrm{rDNA}})}{\sqrt{\left(\mathrm{Var}(\hat{c}(t)_i^{\mathrm{rRNA}}) + \mathrm{Var}(\hat{c}(t)_i^{\mathrm{rDNA}})\right) \cdot 2\mathrm{Var}(\hat{c}(t)_i^{\mathrm{rDNA}})}} && \text{(S33b)} \\
&= \frac{\mathrm{Var}(\hat{c}(t)_i^{\mathrm{rDNA}})}{\sqrt{\left(\mathrm{Var}(\hat{c}(t)_i^{\mathrm{rRNA}}) + \mathrm{Var}(\hat{c}(t)_i^{\mathrm{rDNA}})\right) \cdot 2\mathrm{Var}(\hat{c}(t)_i^{\mathrm{rDNA}})}} && \text{(S33c)}
\end{aligned}$$

which is non-zero as variances are $> 0$ by definition. If rRNA and rDNA have equal variances then the expected correlation reduces to $\frac{1}{2}$. This result is an example of how relying on rRNA:rDNA can introduce spurious correlations, an undesirable statistical artifact.

We can gain greater insight by examining whether there is a *phase delay* between rRNA and rDNA using the cross-power spectral density. This analysis shares some similarities to the phase difference analysis performed using parameters inferred from our gamma + time-varying mean model, though it provides the advantage of allowing us to directly investigate the joint phase relationship over the entire phase spectrum rather than at a single frequency. We start by examining how rRNA:rDNA can covary with rDNA as the time interval $\delta t$ is shifted. This is known as the cross-covariance function

$$R^{(i)}_{\hat{\phi},\,\delta\hat{c}^{\mathrm{rDNA}}}(\delta t) \equiv \overline{\hat{\phi}_i(t) \cdot \delta\hat{c}_i(t+\delta t)^{\mathrm{rDNA}}} = \mathrm{lim}_{T\to\infty} \frac{1}{T} \int_0^T \hat{\phi}_i(t) \cdot \delta\hat{c}_i(t+\delta t)^{\mathrm{rDNA}} dt \qquad \text{(S34)}$$

where the bar represents the average taken *over time* $(t)$. The Fourier transform of this function is the Cross-Power Spectral Density (CPSD), our object of interest

$$\mathcal{S}^{(i)}_{\hat{\phi},\,\delta\hat{c}^{\mathrm{rDNA}}}(f) = \int_{-\infty}^{\infty} R^{(i)}_{\hat{\phi},\,\delta\hat{c}^{\mathrm{rDNA}}}(\delta t)\, e^{-i2\pi f\delta t}\, d\delta t \qquad \text{(S35)}$$

This function captures the power distribution between our variables as signals, allowing for their correlation to be quantified at a given frequency, $f$. While this function is general in that it can be applied to a variety of time series, we note its recent application to the inference of resource competition in microbial community timeseries data (21). To ultimately simplify the CPSD, we can expand the cross-correlation function

$$R^{(i)}_{\hat{\phi},\,\delta\hat{c}^{\mathrm{rDNA}}}(\delta t) = \overline{\left(\hat{c}_i^{\mathrm{rRNA}}(t) - \hat{c}_i^{\mathrm{rDNA}}(t)\right)\left(\hat{c}_i^{\mathrm{rDNA}}(t+\delta t+\Delta t) - \hat{c}_i^{\mathrm{rDNA}}(t+\delta t)\right)}$$

Here $\Delta t$ represents the length of time between *samples*. We then linearly expand the function to obtain

$$R^{(i)}_{\hat{\phi},\,\delta\hat{c}^{\text{rDNA}}}(\delta t) = R^{(i)}_{\hat{c}^{\text{rRNA}},\,\hat{c}^{\text{rDNA}}}(\delta t + \Delta t) - R^{(i)}_{\hat{c}^{\text{rRNA}},\,\hat{c}^{\text{rDNA}}}(\delta t) - R^{(i)}_{\hat{c}^{\text{rDNA}},\,\hat{c}^{\text{rDNA}}}(\delta t + \Delta t) + R^{(i)}_{\hat{c}^{\text{rDNA}},\,\hat{c}^{\text{rDNA}}}(\delta t) \tag{S36}$$

These terms can be consolidated into two interpretable parts: 1) the cross-terms for rRNA and rDNA and 2) the auto-terms of rDNA

$$R^{(i)}_{\hat{\phi},\,\delta\hat{c}^{\text{rDNA}}}(\delta t) = \underbrace{\left[R^{(i)}_{\hat{c}^{\text{rRNA}},\,\hat{c}^{\text{rDNA}}}(\delta t + \Delta t) - R^{(i)}_{\hat{c}^{\text{rRNA}},\,\hat{c}^{\text{rDNA}}}(\delta t)\right]}_{\text{rRNA}-\text{rDNA cross}-\text{terms}} - \underbrace{\left[R^{(i)}_{\hat{c}^{\text{rDNA}},\,\hat{c}^{\text{rDNA}}}(\delta t + \Delta t) - R^{(i)}_{\hat{c}^{\text{rDNA}},\,\hat{c}^{\text{rDNA}}}(\delta t)\right]}_{\text{rDNA auto}-\text{terms}} \tag{S37}$$

The integral for the CPSD is then

$$\mathcal{S}^{(i)}_{\hat{\phi},\,\delta\hat{c}^{\text{rDNA}}}(f) = \int_{-\infty}^{\infty} R^{(i)}_{\hat{c}^{\text{rRNA}},\,\hat{c}^{\text{rDNA}}}(\delta t + \Delta t)\, e^{-i2\pi f\,\delta t}\, d\delta t \;-\; \int_{-\infty}^{\infty} R^{(i)}_{\hat{c}^{\text{rRNA}},\,\hat{c}^{\text{rDNA}}}(\delta t)\, e^{-i2\pi f\,\delta t}\, d\delta t - \int_{-\infty}^{\infty} R^{(i)}_{\hat{c}^{\text{rDNA}},\,\hat{c}^{\text{rDNA}}}(\delta t + \Delta t)\, e^{-i2\pi f\,\delta t}\, d\delta t \;+\; \int_{-\infty}^{\infty} R^{(i)}_{\hat{c}^{\text{rDNA}},\,\hat{c}^{\text{rDNA}}}(\delta t)\, e^{-i2\pi f\,\delta t}\, d\delta t. \tag{S38}$$

We can then use the following shift theorem property

$$\int_{-\infty}^{\infty} R(\delta t + \Delta t)\, e^{-i2\pi f\,\delta t}\, d\delta t \;=\; e^{\,i2\pi f\,\Delta t}\,\mathcal{S}(f). \tag{S39}$$

Which we apply to each shifted integral

$$\begin{aligned}\mathcal{S}^{(i)}_{\hat{\phi},\,\delta\hat{c}^{\text{rDNA}}}(f) &= e^{\,i2\pi f\,\Delta t}\,\mathcal{S}^{(i)}_{\hat{c}^{\text{rRNA}},\,\hat{c}^{\text{rDNA}}}(f) - \mathcal{S}^{(i)}_{\hat{c}^{\text{rRNA}},\,\hat{c}^{\text{rDNA}}}(f) - e^{\,i2\pi f\,\Delta t}\,\mathcal{S}^{(i)}_{\hat{c}^{\text{rDNA}},\,\hat{c}^{\text{rDNA}}}(f) + \mathcal{S}^{(i)}_{\hat{c}^{\text{rDNA}},\,\hat{c}^{\text{rDNA}}}(f)\\ &= \left(e^{\,i2\pi f\,\Delta t} - 1\right)\left[\mathcal{S}^{(i)}_{\hat{c}^{\text{rRNA}},\,\hat{c}^{\text{rDNA}}}(f) - \mathcal{S}^{(i)}_{\hat{c}^{\text{rDNA}},\,\hat{c}^{\text{rDNA}}}(f)\right].\end{aligned} \tag{S40}$$

This result can be interpreted as the difference between the CPSD of rRNA vs. rDNA and that of rDNA vs. rDNA, the latter of which is simply the Power Spectral Density. This difference is then multiplied by a finite-difference transfer function that reduces to $i2\pi f$ as $\Delta t \to 0$. This transfer function is operating as a high-pass filter that removes information for low values of $f$, depends on the choice of $\Delta t$, and the exponential distorts the phase.

Given that our result contains the CPSD between rRNA and rDNA, we choose to examine the statistical dependency between these two variables. Specifically, we focus on their lead/lag behavior. To determine the time lead/lag between rRNA and rDNA, we then convert the CPSD to the radian domain by calculating the phase angle, which represents the phase difference between rRNA and rDNA at a given value of $f$.

$$\psi^{(i)}_{\text{rRNA, rDNA}}(f) = \arg(\mathcal{S}^{(i)}_{\text{rRNA, rDNA}}(f)) \tag{S41}$$

from which we obtain the time lag

$$\mathcal{T}^{(i)}_{\text{rRNA, rDNA}}(f) = \frac{\psi^{(i)}_{\text{rRNA, rDNA}}(f)}{2\pi f} \tag{S42}$$

Due to the limitations of sampling, it is desirable to remove frequencies that likely represent noise rather than signal. We can identify said frequencies using the magnitude-squared coherence, a measure of the fraction of power at a given frequency that is shared between our two timeseries.

$$\mathrm{MSC}^{(i)}_{\mathrm{rRNA,\,rDNA}}(f) = \frac{|\mathcal{S}^{(i)}_{\mathrm{rRNA,\,rDNA}}(f)|^2}{\mathcal{S}^{(i)}_{\mathrm{rRNA,\,rRNA}}(f) \cdot \mathcal{S}^{(i)}_{\mathrm{rDNA,\,rDNA}}(f)}. \tag{S43}$$

From which we extract values of $f$ where the coherence exceeds a specified minimum, $\mathcal{F}_\rangle = \{\, f \mid \mathrm{MSC}^{(i)}_{\mathrm{rRNA,\,rDNA}}(f) > \mathrm{MSC}_{\mathrm{min}} \,\}$. We then arrive at our definition of the cross-spectral time shift.

$$\overline{\mathcal{T}^{(i)}}_{\mathrm{rRNA,\,rDNA}} = \frac{\sum_{f \in \mathcal{F}_i} \mathcal{T}^{(i)}_{\mathrm{rRNA,\,rDNA}}(f) \cdot \mathrm{MSC}^{(i)}(f)}{\sum_{f \in \mathcal{F}_i} \mathrm{MSC}^{(i)}(f)} \tag{S44}$$

To identify a null distribution of $\overline{\mathcal{T}^{(i)}}_{\mathrm{rRNA,\,rDNA}}$ in the absence of underlying lead or lag behavior, we generated signals where the phase of the data is randomized by the spectrum amplitude is retained, known as surrogate signals (22). Because we are using the fast Fourier transform (FFT) when performing the above calculations, we will discuss the null in terms of discrete signals. First,

$$C^{(i)}_{\mathrm{rRNA}}(k) = \sum_{m=0}^{M-1} \hat{c}^{\mathrm{rRNA}}_{i,m} e^{-i2\pi k \frac{m}{M}} \tag{S45}$$

where $k = 0, ..., M-1$ is the frequency index. we then generate random phases $\psi^{(s)} \sim \mathrm{Uniform}[0, 2\pi)$ for $1 \leq k \leq M/2 - 1$. Surrogate Fourier coefficients are calculated as follows

$$(C^{(i)}_{\mathrm{rRNA}}(k))^{(s)} = \begin{cases} |C^{(i)}_{\mathrm{rRNA}}(0)|, & k = 0 \quad \text{(DC component, unchanged)} \\ |C^{(i)}_{\mathrm{rRNA}}(k)| e^{i\psi^{(s)}(k)}, & 1 \leq k \leq M/2 - 1 \\ |C^{(i)}_{\mathrm{rRNA}}(\frac{M}{2})|, & k = \frac{M}{2} \\ (C^{(i)}_{\mathrm{rRNA}}(M-k))^{s,*}, & M/2 + 1 \leq k \leq M - 1 \end{cases}$$

where the last rule uses the complex conjugate of the positive frequency bin in order to ensure that the inverse FFT remains real-valued. We then perform the inverse FFT to obtain our surrogate rRNA abundances back in the time domain.

$$(\hat{c}^{\mathrm{rRNA}}_{i,m})^{(s)} = \frac{1}{M} \sum_{m=0}^{M-1} (C^{(i)}_{\mathrm{rRNA}}(k))^{(s)} e^{i2\pi k \frac{m}{M}} \tag{S46}$$

And calculated the the mean time lag using the CPSD with the empirical rDNA timeseries and rRNA surrogate timeseries. We chose $\mathrm{MSC}_{\mathrm{min}} = 0.3$ and subsetted frequencies to those $> \frac{1}{\mathrm{median}(\delta t)} \cdot \frac{1}{\#\,\mathrm{samples}}$. Null distributions were obtained using $10^3$ iterations. The CPSD was calculated using the Welch method. Analyses were performed using `SciPy v1.16.0`. For each ASV a $P$-value was calculated using the null. We then performed a *global* test by asking whether there was a significant shift *across all ASVs.* We first calculate a global shift by averaging mean lag over our ASVs

$$\overline{\mathcal{T}}_{\text{global}} = \frac{1}{|S^*|} \sum_{i \in S^*} \overline{\mathcal{T}^{(i)}}_{\text{rRNA, rDNA}} \tag{S47}$$

We then calculate the same statistic for each random iteration $s$

$$\overline{\mathcal{T}^{(s)}}_{\text{global}} = \frac{1}{|S^*|} \sum_{i \in S^*} \overline{\mathcal{T}^{(i,s)}}_{\text{rRNA, rDNA}}. \tag{S48}$$

And obtain a $P$-value using our null distribution.

##### Interpreting empirical rRNA:rDNA patterns through a minimal, physiologically-motivated model

We have two empirical observations from rRNA that require interpretation: 1) time-averaged rRNA:rDNA increases with the number of 16S rRNA operon copies harbored within an ASV and 2) the mean cross-spectral phase delay indicates that rRNA dynamics lead those of rDNA. The first observation rules out a model where the rate of transcription remains constant. The second observation requires feedback between rRNA and rDNA, such as ribosome-limited growth. These two observations provide the motivation to go beyond modeling rRNA and rDNA as separate Langevin equations by considering their physiological interactions.

We start by defining the deterministic dynamics for the total number of cells, 16S rRNA gene copies, and ribosomes within a given ASV. Namely, we are modeling quantities at the level of *populations*. We assume that the change in gene copies over time for a given ASV is proportional to the change in the number of cells multiplied by the number of 16S rRNA operons per-cell ($g$), $\frac{dN_{\text{gene}}}{dt} = g \cdot \frac{dN_{\text{cells}}}{dt}$. We define $\kappa_t \frac{N_{\text{ribo}}}{N_{\text{gene}}}$ as the per-capita birth rate and $bN_{\text{cell}} + \delta$ as a separate per-capita turnover rate representing intraspecific competition and death. The rate of per-capita ribosome assembly and degradation is $k_0 \cdot g^{\alpha+1}$ and $\gamma$, respectively, where $\alpha$ represents the gene dosage effect and $k_0$ is a conversion rate. Together, we obtain the following per-capita reactions:

$$N_{\text{cell}} \xrightarrow{\kappa_t \frac{N_{\text{ribo}}}{N_{\text{gene}}}} 2N_{\text{cell}} \quad \text{(Density-independent cellular division)} \tag{S49a}$$

$$N_{\text{cell}} \xrightarrow{b\,N_{\text{cell}} + \delta} \emptyset \quad \text{(Intraspecific competition + death)} \tag{S49b}$$

$$N_{\text{cell}} \xrightarrow{k_0\,\kappa_t\,g^{\alpha+1}} N_{\text{cell}} + N_{\text{ribo}} \quad \text{(Ribosome assembly)} \tag{S49c}$$

$$N_{\text{ribo}} \xrightarrow{\gamma} \emptyset \quad \text{(Ribosome degradation)} \tag{S49d}$$

Typically at this stage a master equation of the reaction-kinetics are expanded and converted to obtain a Langevin equation. However, this procedure results in fluctuations being driven by demographic noise, whereas we are primarily interested in *environmental* noise. To incorporate this noise source we first translate the above reactions to the following *deterministic* dynamics:

$$\frac{dN_{\text{cell}}}{dt} = \kappa_t \frac{N_{\text{ribo}}}{N_{\text{gene}}} N_{\text{cell}} - b\, N_{\text{cell}}^2 - \delta\, N_{\text{cell}} \tag{S50a}$$

$$\frac{dN_{\text{gene}}}{dt} = g \cdot \frac{dN_{\text{cell}}}{dt} \tag{S50b}$$

$$\frac{dN_{\text{ribo}}}{dt} = k_0\, \kappa_t\, g^{\alpha+1}\, N_{\text{cell}} - \gamma N_{\text{ribo}} \tag{S50c}$$

The ODE for $N_{\text{cell}}$ is similar in form to the deterministic form of the SLM, where $b \equiv \kappa_t \bar{\rho}/K$ recovers the original carrying capacity $K$ when $\delta = 0$. Throughout this analysis we fix the turnover rate $bN_{\text{cell}} + \delta$ at its mean value, reflecting a fixed resource ceiling rather than the rapid metabolic state that drives $\kappa_t$, $k_0$, and $\gamma$. Our justification for this choice is that shared fluctuations between rRNA and rDNA likely occur over timescales much shorter than the seasonal oscillations observed in our system.

We can now add fluctuations in the *rates* as a type of environmental noise. We consider each of our three rates ($\kappa_t$, $k_0$, $\gamma$) as $\lambda_\mu$, fluctuating around its mean value as:

$$\lambda_\mu(t) = \bar{\lambda}_\mu \big[1 + \sigma_\mu \eta_\mu(t)\big], \qquad \langle \eta_\mu(t) \rangle = 0. \tag{S51}$$

resulting in every reaction rate acquiring a multiplicative noise term. The system of Langevin equations can then be obtained by replacing each rate constant with this function. Because all three rate constants respond to a shared environment, it is worth incorporating correlated fluctuations at this stage.

$$\langle \eta_\mu(t)\, \eta_\nu(t') \rangle = C_{\mu\nu}\, \delta(t - t'), \qquad \mathbf{C} = \begin{pmatrix} 1 & \rho_{12} & \rho_{13} \\ \rho_{12} & 1 & \rho_{23} \\ \rho_{13} & \rho_{23} & 1 \end{pmatrix}, \tag{S52}$$

with $\mathbf{C}$ being positive semi-definite. These coupling parameter can be interpreted. For example, $\rho_{23}$ represents the coupling $k_0 \leftrightarrow \gamma$, which could be driven by environment-induced fluctuations in RNA-polymerase simultaneously altering rates of ribosome assembly and degradation.

Applying Eq. S51 to each reaction in our deterministic system of ODEs, we obtain our system of Langevins

$$\frac{dN_{\text{cell}}}{dt} = \kappa_t \frac{N_{\text{ribo}}}{N_{\text{gene}}} N_{\text{cell}}\big(1 + \sigma_1 \eta_1(t)\big) - b\, N_{\text{cell}}^2 - \delta\, N_{\text{cell}}, \tag{S53a}$$

$$\frac{dN_{\text{gene}}}{dt} = g\, \frac{dN_{\text{cell}}}{dt}, \tag{S53b}$$

$$\frac{dN_{\text{ribo}}}{dt} = k_0\, \kappa_t\, g^{\alpha+1} N_{\text{cell}}\big(1 + \sigma_2 \eta_2(t)\big) - \gamma N_{\text{ribo}}\big(1 + \sigma_3 \eta_3(t)\big). \tag{S53c}$$

**Deriving $g$ vs. steady-state rRNA:rDNA**

Setting all noise terms to zero and using the exact identity $\kappa_t(N_{\text{ribo}}/N_{\text{gene}})N_{\text{cell}} = (\kappa_t/g)N_{\text{ribo}}$, the $N_{\text{cell}}$ steady state satisfies

$$\frac{\kappa_t}{g} \bar{N}_{\text{ribo}} = b\, \bar{N}_{\text{cell}}^2 + \delta\, \bar{N}_{\text{cell}}. \tag{S54}$$

Setting $dN_{\text{ribo}}/dt = 0$, with the ribosome assembly rate now coupled to translational capacity, yields

$$k_0\,\kappa_t\,g^{\alpha+1}\,\bar{N}_{\text{cell}} = \gamma\,N^*_{\text{ribo}}, \qquad N^*_{\text{ribo}} = \frac{k_0\,\kappa_t\,g^{\alpha+1}\,\bar{N}_{\text{cell}}}{\gamma}. \tag{S55}$$

Substituting Eq. S55 into Eq. S54 and solving gives $\bar{N}_{\text{cell}} = (\alpha_r - \delta)/b$, with $\alpha_r \equiv \kappa_t\bar{\rho}$, $\bar{\rho} \equiv \bar{N}_{\text{ribo}}/\bar{N}_{\text{gene}}$, equivalently $\bar{N}_{\text{cell}} = K(1 - \delta/\alpha_r) < K$, strictly below the carrying capacity rather than sitting exactly at it. $N^*_{\text{gene}} = g\bar{N}_{\text{cell}}$. Dividing Eq. S55 by $N^*_{\text{gene}}$,

$$\frac{N^*_{\text{ribo}}}{N^*_{\text{gene}}} = \frac{k_0\,\kappa_t\,g^{\alpha+1}\,\bar{N}_{\text{cell}}/\gamma}{g\,\bar{N}_{\text{cell}}} = \frac{k_0}{\gamma}\,\kappa_t\,g^{\alpha}, \tag{S56}$$

so that

$$\ln\!\left(\frac{N^*_{\text{ribo}}}{N^*_{\text{gene}}}\right) = \ln\frac{k_0}{\gamma} + \ln\kappa_t + \alpha\ln g, \tag{S57}$$

$\bar{N}_{\text{cell}}$ cancels in the ratio, so the log-linear relationship and its slope $\alpha$ in $\ln g$ are insensitive to intraspecific competition and death rate. Substituting $\bar{\rho} = (k_0/\gamma)\kappa_t g^{\alpha}$ back into the definition of $\alpha_r$ gives the growth rate parameter in terms of the rates,

$$\alpha_r = \kappa_t\bar{\rho} = \frac{k_0}{\gamma}\,\kappa_t^2\,g^{\alpha}, \tag{S58}$$

which can be inverted and substituted back to express the steady-state ratio directly in terms of the growth parameters,

$$\bar{\rho} = \sqrt{\frac{k_0\,g^{\alpha}}{\gamma}\,\alpha_r}. \tag{S59}$$

Equation S59 shows that the steady-state rRNA:rDNA ratio increases monotonically with the growth rate parameter $\alpha_r$ and log-linearly with operon copy number $g$ (Eq. S57).

**rRNA vs. rDNA cross-spectral phase delay**

As we are interested in the logarithm of our variables, we start with the log-linearization around the steady-state $(\bar{N}_{\text{cell}}, \bar{N}_{\text{ribo}})$:

$$\delta n = \ln N_{\text{cell}} - \ln\bar{N}_{\text{cell}}, \qquad \delta r = \ln N_{\text{ribo}} - \ln\bar{N}_{\text{ribo}}. \tag{S60}$$

Linearizing around $(\bar{N}_{\text{cell}}, \bar{N}_{\text{ribo}})$ and dividing by $\bar{N}_{\text{cell}}$ gives the coefficients

$$\lambda_n = 2\alpha_r - \delta, \qquad \alpha_r = \kappa_t\bar{\rho}, \qquad \tilde{\sigma}_n = \alpha_r\,\sigma_1, \tag{S61}$$

using $b\bar{N}_{\text{cell}} = \alpha_r - \delta$ at the fixed point. For $\delta r$, we denote the stochastic parameters

$$\tilde{\sigma}_+ = \sigma_2\gamma, \qquad \tilde{\sigma}_- = \sigma_3\gamma. \tag{S62}$$

Using these terms, the linearized system is

$$\frac{d(\delta n)}{dt} = -\lambda_n\,\delta n + \alpha_r\,\delta r + \tilde{\sigma}_n\,\eta_1(t), \tag{S63a}$$

$$\frac{d(\delta r)}{dt} = \gamma(\delta n - \delta r) + \tilde{\sigma}_+\,\eta_2(t) - \tilde{\sigma}_-\,\eta_3(t), \tag{S63b}$$

with the following three effective noise amplitudes

$$\tilde{\sigma}_n = \alpha_r\,\sigma_1, \qquad \tilde{\sigma}_+ = \sigma_2\gamma, \qquad \tilde{\sigma}_- = \sigma_3\gamma. \tag{S64}$$

The Fourier transform of our system is then

$$\begin{pmatrix} i\omega + \lambda_n & -\alpha_r \\ -\gamma & i\omega + \gamma \end{pmatrix} \begin{pmatrix} \delta\hat{n} \\ \delta\hat{r} \end{pmatrix} = \begin{pmatrix} \tilde{\sigma}_n\hat{\eta}_1 \\ \tilde{\sigma}_+\hat{\eta}_2 - \tilde{\sigma}_-\hat{\eta}_3 \end{pmatrix}, \tag{S65}$$

which has the characteristic polynomial

$$D(\omega) = (i\omega + \lambda_n)(i\omega + \gamma) - \alpha_r\gamma. \tag{S66}$$

Using Cramer's rule, we obtain a solution in terms of the determinants of our matrix:

$$D\,\delta\hat{n} = \underbrace{(i\omega + \gamma)\tilde{\sigma}_n}_{B_1}\,\hat{\eta}_1 + \underbrace{\alpha_r\tilde{\sigma}_+}_{B_2}\,\hat{\eta}_2 + \underbrace{-\alpha_r\tilde{\sigma}_-}_{B_3}\,\hat{\eta}_3, \tag{S67a}$$

$$D\,\delta\hat{r} = \underbrace{\gamma\tilde{\sigma}_n}_{A_1}\,\hat{\eta}_1 + \underbrace{(i\omega + \lambda_n)\tilde{\sigma}_+}_{A_2}\,\hat{\eta}_2 + \underbrace{-(i\omega + \lambda_n)\tilde{\sigma}_-}_{A_3}\,\hat{\eta}_3. \tag{S67b}$$

We can now derive the CPSD. Using $\langle\hat{\eta}_\mu\hat{\eta}_\nu^*\rangle = C_{\mu\nu}$, we obtain the following general expression:

$$|D|^2\,\mathcal{S}_{rn}(\omega) = \mathbb{E}[D\delta\hat{r}\cdot(D\delta\hat{n})^*] = \sum_{\mu,\nu=1}^{3} A_\mu B_\nu^* C_{\mu\nu}, \tag{S68}$$

with the diagonal terms ($C_{\mu\mu} = 1$)

$$\begin{aligned} A_1B_1^* &= \gamma\tilde{\sigma}_n^2(-i\omega + \gamma), \\ A_2B_2^* &= \alpha_r\tilde{\sigma}_+^2(i\omega + \lambda_n), \\ A_3B_3^* &= \alpha_r\tilde{\sigma}_-^2(i\omega + \lambda_n). \end{aligned}$$

Defining the total rRNA noise variance

$$\tilde{\Sigma}_r^2 \equiv \tilde{\sigma}_+^2 + \tilde{\sigma}_-^2 = \gamma^2(\sigma_2^2 + \sigma_3^2), \tag{S69}$$

we obtain the diagonal sum

$$\gamma\tilde{\sigma}_n^2(-i\omega + \gamma) + \alpha_r\tilde{\Sigma}_r^2(i\omega + \lambda_n). \tag{S70}$$

The off-diagonal terms can be reduced by first defining

$$\Phi(\omega) \equiv \gamma(\alpha_r + \lambda_n) + \omega^2 + i\omega(\gamma - \lambda_n), \tag{S71}$$

which we have obtained by expanding the factored form.

$$A_1 B_2^* + A_2 B_1^* = \rho_{12}\tilde{\sigma}_n \tilde{\sigma}_+ \, \Phi(\omega), \tag{S72}$$
$$A_1 B_3^* + A_3 B_1^* = -\rho_{13}\tilde{\sigma}_n \tilde{\sigma}_- \, \Phi(\omega), \tag{S73}$$
$$A_2 B_3^* + A_3 B_2^* = -2\rho_{23}\alpha_r (i\omega + \lambda_n)\tilde{\sigma}_+ \tilde{\sigma}_-. \tag{S74}$$

Using these diagonal and off-diagonal terms, we obtain a full expression of the CPSD:

$$|D|^2 \mathcal{S}_{rn} = \gamma\tilde{\sigma}_n^2(-i\omega + \gamma) + \alpha_r \tilde{\Sigma}_r^2 (i\omega + \lambda_n) + \tilde{\sigma}_n(\rho_{12}\tilde{\sigma}_+ - \rho_{13}\tilde{\sigma}_-)\Phi(\omega) - 2\rho_{23}\alpha_r \tilde{\sigma}_+ \tilde{\sigma}_- (i\omega + \lambda_n). \tag{S75}$$

We can obtain the imaginary and real parts using $\mathrm{Im}[\Phi] = \omega(\gamma - \lambda_n)$ and $\mathrm{Re}[\Phi] = \gamma(\alpha_r + \lambda_n) + \omega^2$:

$$\mathrm{Im}[\mathcal{S}_{rn}] = \frac{\omega \mathcal{N}}{|D(\omega)|^2}, \tag{S76a}$$

$$\mathrm{Re}[\mathcal{S}_{rn}] = \frac{\gamma^2\tilde{\sigma}_n^2 + \alpha_r\lambda_n(\tilde{\Sigma}_r^2 - 2\rho_{23}\tilde{\sigma}_+\tilde{\sigma}_-) + \tilde{\sigma}_n(\rho_{12}\tilde{\sigma}_+ - \rho_{13}\tilde{\sigma}_-)[\gamma(\alpha_r + \lambda_n) + \omega^2]}{|D(\omega)|^2}, \tag{S76b}$$

where we have defined

$$\mathcal{N} \equiv \underbrace{\alpha_r\tilde{\Sigma}_r^2 - \gamma\tilde{\sigma}_n^2}_{\text{uncoupled}} + \underbrace{(\gamma - \lambda_n)\tilde{\sigma}_n(\rho_{12}\tilde{\sigma}_+ - \rho_{13}\tilde{\sigma}_-) - 2\rho_{23}\alpha_r\tilde{\sigma}_+\tilde{\sigma}_-}_{\text{coupling corrections}}. \tag{S77}$$

The absence of $\omega$ inside $\mathcal{N}$ means that the direction of the phase lead does not depend on frequency. The real part is frequency-dependent through the $\omega^2$ term in the coupling corrections, a behavior that is absent when $\rho_{12} = 0$ or $\rho_{13} = 0$.

We can now identify the lead criterion for rRNA. Here, $\psi_{rn} > 0$ means $\ln N_{\text{ribo}}$ *leads* $\ln N_{\text{gene}}$:

$$\begin{aligned}\psi_{rn}(\omega) &= \arg(\mathcal{S}_{rn}(\omega)) \\ &= \arctan\left(\frac{\mathrm{Im}[\mathcal{S}_{rn}(\omega)]}{\mathrm{Re}[\mathcal{S}_{rn}(\omega)]}\right) \\ &= \arctan\left(\frac{\omega\mathcal{N}}{\gamma^2\tilde{\sigma}_n^2 + \alpha_r\lambda_n(\tilde{\Sigma}_r^2 - 2\rho_{23}\tilde{\sigma}_+\tilde{\sigma}_-) + \tilde{\sigma}_n(\rho_{12}\tilde{\sigma}_+ - \rho_{13}\tilde{\sigma}_-)[\gamma(\alpha_r + \lambda_n) + \omega^2]}\right).\end{aligned} \tag{S78}$$

The predicted time lag is $\mathcal{T}(\omega) = \psi_{rn}(\omega)/\omega$. In the small-angle limit $\arctan(x) \approx x$ and dropping the $\omega^2$ term:

$$\mathcal{T} \approx \frac{\mathcal{N}}{\gamma^2\tilde{\sigma}_n^2 + \alpha_r\lambda_n(\tilde{\Sigma}_r^2 - 2\rho_{23}\tilde{\sigma}_+\tilde{\sigma}_-) + \tilde{\sigma}_n(\rho_{12}\tilde{\sigma}_+ - \rho_{13}\tilde{\sigma}_-)\gamma(\alpha_r + \lambda_n)}. \tag{S79}$$

So $\psi_{rn} > 0$ if and only if $\mathcal{N} > 0$

$$\alpha_r\tilde{\Sigma}_r^2 - \gamma\tilde{\sigma}_n^2 + (\gamma - \lambda_n)\tilde{\sigma}_n(\rho_{12}\tilde{\sigma}_+ - \rho_{13}\tilde{\sigma}_-) - 2\rho_{23}\alpha_r\tilde{\sigma}_+\tilde{\sigma}_- > 0. \tag{S80}$$

Given the number of terms in this expression, it is useful to identify limiting cases with biological meaning. We first consider the scenario where all noise sources are uncoupled, $\rho_{\mu\mu'} = 0$. The criterion reduces to

$$\frac{\alpha_r}{\gamma} > \frac{\tilde{\sigma}_n^2}{\tilde{\Sigma}_r^2} = \frac{\sigma_1^2\alpha_r^2}{\gamma^2(\sigma_2^2 + \sigma_3^2)}, \tag{S81}$$

which can be interpreted as rRNA *leading* rDNA when the ratio of the ribosome assembly and degradation rates exceeds the ratio of the strength of cellular division noise to ribosomal noise. In the limit where cell division is deterministic ($\tilde{\sigma}_n \to 0$), rRNA will lead rDNA if there is any ribosomal noise ($\tilde{\Sigma}_r^2 > 0$). In contrast, if ribosomal dynamics are deterministic ($\tilde{\Sigma}_r^2 = 0$) and noise is driven solely by cellular division ($\tilde{\sigma}_n^2 > 0$), then rRNA *always* lags rDNA, driven passively by cell-division noise.

In the main manuscript we present the system of Langevins in this parameter limit, setting $\sigma_2 = \sigma_3$ and renaming the subscripts to obtain

$$\frac{dN_{\text{cell}}}{dt} = \kappa_t\,\frac{N_{\text{ribo}}}{N_{\text{gene}}}\,N_{\text{cell}}\big(1 + \sigma_{\text{cell}}\eta_{\text{cell}}(t)\big) - N_{\text{cell}}^2 - N_{\text{cell}}, \tag{S82a}$$

$$\frac{dN_{\text{gene}}}{dt} = g\,\frac{dN_{\text{cell}}}{dt}, \tag{S82b}$$

$$\frac{dN_{\text{ribo}}}{dt} = k_0\,\kappa_t\,g^{\alpha+1}N_{\text{cell}} - \gamma N_{\text{ribo}} + \sigma_{\text{ribo}}\sqrt{k_0^2\,\kappa_t^2\,g^{2(\alpha+1)}N_{\text{cell}}^2 + \gamma^2 N_{\text{ribo}}^2}\,\eta_{\text{ribo}}(t), \tag{S82c}$$

where we applied the law of total variance for independent sources of noise to obtain a single effective noise term for ribosomes and set $b = \delta = 1$. We briefly note that rRNA:rDNA is then

$$\begin{aligned}\log\left(\frac{N_{\text{ribo}}(t)}{N_{\text{gene}}(t)}\right) &= \log\left[k_0\,\kappa_t\,g^{\alpha+1}N_{\text{cell}} - \gamma N_{\text{ribo}} + \sigma_{\text{ribo}}\sqrt{k_0^2\,\kappa_t^2\,g^{2(\alpha+1)}N_{\text{cell}}^2 + \gamma^2 N_{\text{ribo}}^2}\,\eta_{\text{ribo}}(t)\right] - \log\big(N_{\text{ribo}}\big)\\ &\quad - \log\left[\kappa_t\,\frac{N_{\text{ribo}}}{N_{\text{gene}}}\big(1 + \sigma_{\text{cell}}\eta_{\text{cell}}(t)\big) - N_{\text{cell}} - 1\right]\end{aligned} \tag{S83}$$

Returning to our CPSD results, the lead criterion reduces to

$$\frac{\alpha_r}{\gamma} > \frac{\sigma_{\text{cell}}^2\,\alpha_r^2}{2\gamma^2\,\sigma_{\text{ribo}}^2} \iff \sigma_{\text{ribo}}^2 > \frac{\sigma_{\text{cell}}^2\,\alpha_r}{2\gamma}. \tag{S84}$$

The term $\alpha_r$ (Eq. S58) grows quadratically with $\kappa_t$, so the right-hand side of Eq. S84 increases with growth rate. Faster-growing populations therefore require proportionally larger ribosomal noise amplitude $\sigma_{\text{ribo}}^2$ for rRNA to lead rDNA, linking the model's growth-rate dependence (Eq. S59) to its observed behavior.

Similarly, we can examine the opposite limit where all noise sources are fully coupled, $\rho_{\mu\mu'} = 1$. The criterion for $\mathcal{N} > 0$ can be obtained by substituting Eq. S61 and dividing by $\alpha_r\gamma$,

$$\gamma(\sigma_2 - \sigma_3)^2 + (\gamma - \lambda_n)\,\sigma_1(\sigma_2 - \sigma_3) > \alpha_r\sigma_1^2, \qquad \lambda_n = 2\alpha_r - \delta = 2\alpha_r - 1. \tag{S85}$$

There are two regimes: 1) $\sigma_2 = \sigma_3$, for which the left-hand side vanishes and the inequality is never satisfied, where rRNA always lags rDNA and 2) $\sigma_2 \neq \sigma_3$, for which the criterion can be written as

$$\frac{\tilde{\sigma}_n}{\tilde{\sigma}_+ - \tilde{\sigma}_-}\left(\frac{\gamma\tilde{\sigma}_n}{\tilde{\sigma}_+ - \tilde{\sigma}_-} - (\gamma - \lambda_n)\right) < \alpha_r, \tag{S86}$$

**Recapitulating stationary gamma AFDs**

We derived the stationary distribution for our coupled model of cellular and ribosomal growth to see whether it is qualitatively consistent with our earlier, phenomenological model of a time-varying mean with gamma-distributed fluctuations. Namely, we are trying to recapitulate a *stationary* gamma distribution for $N_{\text{cell}}$ and $N_{\text{ribo}}$. For an Itô Langevin equation of form

$$\frac{dx}{dt} = f(x) + h(x)\,\eta(t), \tag{S87}$$

the corresponding Fokker–Planck equation is $\partial_t P = -\partial_x(fP) + \frac{1}{2}\partial_x^2(h^2 P)$. Setting the time derivative to zero and integrating, one obtains

$$P^*(x) \propto \frac{1}{h(x)^2}\,\exp\left[2\int^x \frac{f(x')}{h(x')^2}\,dx'\right]. \tag{S88}$$

Plugging in the terms for the $N_{\text{cell}}$ Langevin, we obtain

$$P(N_{\text{cell}}) \propto N_{\text{cell}}^{\,2(\alpha_r-\delta)/(\sigma_{\text{cell}}^2\alpha_r^2)\,-\,2}\,\exp\left[-\frac{2b}{\sigma_{\text{cell}}^2\alpha_r^2}N_{\text{cell}}\right], \tag{S89}$$

which is the Gamma distribution.

For $N_{\text{ribo}}$, we apply the mean-field treatment to $N_{\text{cell}} \approx \bar{N}_{\text{cell}}$ in Eq. S82c to obtain $f(N_{\text{ribo}}) = A - \gamma N_{\text{ribo}}$, $g(N_{\text{ribo}}) = \sigma_{\text{ribo}}\sqrt{A^2 + \gamma^2 N_{\text{ribo}}^2}$, $A \equiv k_0\kappa_t g^{\alpha+1}\bar{N}_{\text{cell}}$. The integral can be evaluated by splitting the integrand into two pieces

$$\frac{f(N_{\text{ribo}})}{g(N_{\text{ribo}})^2} = \frac{A - \gamma N_{\text{ribo}}}{\sigma_{\text{ribo}}^2(A^2 + \gamma^2 N_{\text{ribo}}^2)} = \frac{1}{\sigma_{\text{ribo}}^2}\left[\frac{A}{A^2 + \gamma^2 N_{\text{ribo}}^2} - \frac{\gamma N_{\text{ribo}}}{A^2 + \gamma^2 N_{\text{ribo}}^2}\right]. \tag{S90}$$

We recognize that the integral of the first term as the inverse tangent function and that the second is logarithmic, returning

$$2\int \frac{f(N'_{\text{ribo}})}{g(N'_{\text{ribo}})^2}\,dN'_{\text{ribo}} = \frac{2}{\sigma_{\text{ribo}}^2\gamma}\arctan\left(\frac{\gamma N_{\text{ribo}}}{A}\right) - \frac{1}{\sigma_{\text{ribo}}^2\gamma}\ln\left(A^2 + \gamma^2 N_{\text{ribo}}^2\right), \tag{S91}$$

where we ignore the arbitrary constant of integration as it does not affect the identity of the stationary distribution. From this result, we obtain a distribution known as the Pearson type IV distribution

$$P(N_{\text{ribo}}) \propto \left(A^2 + \gamma^2 N_{\text{ribo}}^2\right)^{-1-\frac{1}{\sigma_{\text{ribo}}^2\gamma}}\exp\left[\frac{2}{\sigma_{\text{ribo}}^2\gamma}\arctan\left(\frac{\gamma N_{\text{ribo}}}{A}\right)\right]. \tag{S92}$$

This is not a gamma distribution, as it interpolates between Gaussian behavior near $N_{\text{ribo}} = 0$ and an inverse-gamma in the tail. However, this distribution is qualitatively similar to a gamma distribution when the strength of noise is not too strong. Therefore, our choice of a coupled model of rRNA–rDNA dynamics is consistent with the form of the empirical probability distributions of both rRNA and rDNA, observations that served as the motivation for our initial uncoupled model (i.e., time-varying mean with gamma fluctuations).

#### Sensitivity analysis

For each ASV $i$, a single joint Generalized Additive Model (GAM) was fitted with all three response variables (rDNA, rRNA, rRNA:rDNA).

$$y_{i,g,t} = \mu_k + \sum_{j=1}^{J} f_{j,k,i}(x_{j,t}) + \varepsilon_{i,k,t}, \qquad \varepsilon_{i,k,t} \sim \mathcal{N}(0, \sigma^2), \tag{S93}$$

where $x_{i,g,t}$ is the value of data type $k \in \{\text{rDNA, rRNA, rRNA:rDNA}\}$ at time $t$. Here $\mu_k$ is a data-type-specific intercept, and $f_{j,k,i}(z_{j,t})$ is the smooth partial effect of predictor $j$ at value $z_{j,t}$, estimated separately for each environmental variable as a factor interaction. The hat notation $\hat{f}_{j,k,i}$ denotes the estimated smooth from the fitted model and $\varepsilon_{i,k,t}$ represents zero-centered Gaussian error.

Model fitting is performed for all three predictors so that a simultaneous confidence interval of the sensitivity can be calculated from the joint posterior covariance matrix. The nine environmental predictors were each fitted with thin-plate regression splines with $k = 5$ basis functions. The cyclic temporal predictor (day of year) used a cyclic cubic regression spline with $k = 12$, and the non-cyclic temporal trend (days since the start of the study) used a thin-plate spline with $k = 10$. All models were fitted with `mgcv::gam` using `method = ‘‘REML‘‘` (23).

Two complementary diagnostics were run for each of the 21 ASV GAMs using the `extract_diagnostics` function. 1) The adequacy of the basis dimension $k$ was assessed with `gratia::k_check` by computing a randomization test to determine whether the residuals of each smooth displayed significant autocorrelation (23). A smooth was flagged as having failed if both $P < 0.05$ and the $k$-index $< 1$. No smooth terms failed the $k$-check across all 21 joint GAMs. 2) The fraction of null deviance explained by the model was examined for each GAM as a global measure of fit. A range of 0.417 - 0.886 was found across ASVs with a mean of 0.72. We elected to retain all 21 joint GAMs for the sensitivity analysis.

##### Sensitivity index

We estimated the global sensitivity of sequence data type $k$ to environmental variable $j$ for each ASV as the mean absolute value of the first derivative of the estimated smooth, evaluated on a regular grid of $M$ points that spans the observed range of $z_j$.

$$I^{\text{GAM}}_{j,k,i} = \frac{1}{M} \sum_{m=1}^{M} \left| \frac{d\hat{f}_{j,k,i}}{dz_j} \Big|_{z_j = z_m} \right|, \tag{S94}$$

where $\{z_1, \ldots, z_M\}$ is a regular evaluation grid of points spanning the observed range of predictor $j$. Derivatives were estimated by central finite differences with simultaneous 95% confidence intervals via the `gratia` (24). Here $I^{\text{GAM}}_{j,k,i}$ is interpreted as the average rate of change of the modeled response per-unit increase in environmental variable $j$ across its observed range.

To test whether rRNA:rDNA is more sensitive than rDNA across, we computed the difference in integrated sensitivity for each ASV:

$$d^{(\text{alt})}_{j,i} = I^{\text{GAM}}_{j,\text{alt},i} - I^{\text{GAM}}_{j,\text{rDNA},i} \tag{S95}$$

and summarized the effect across $|S^*| = 21$ ASVs as the median scaled difference:

$$\tilde{d}^{(\text{alt})}_{j} = \frac{\text{median}_i \left( d^{(\text{alt})}_{j,i} \right)}{\text{IQR}_i \left( I^{\text{GAM}}_{j,\text{rDNA},i} \right)} \tag{S96}$$

The IQR denominator makes the statistic dimensionless and comparable across predictors with different units and scales, representing the difference between quartiles at 0.75 and 0.25. $\tilde{d}_j > 0$ indicates that the alternative variable is on average more sensitive than rDNA; $\tilde{d}_j < 0$ indicates the opposite. Bootstrap 95% confidence intervals were computed by resampling the 21 ASVs with replacement for $10^4$ iterations. Statistical significance was assessed by a two-sided paired Wilcoxon signed-rank test, with $P$-values corrected by the Benjamini-Hochberg false discovery rate across all comparisons.

We briefly note that $\tilde{d}_j^{\mathrm{rRNA}}$ is not guaranteed to be equal to $\tilde{d}_j^{\mathrm{rRNA:rDNA}}$ because each of these terms represents the absolute value of the difference. Thus, under the triangle inequality the following relation holds

$$\left| \left.\frac{d\hat{f}_{j,\mathrm{rRNA},i}}{dz_j}\right|_{z_j=z_m} - \left.\frac{d\hat{f}_{j,\mathrm{rDNA},i}}{dz_j}\right|_{z_j=z_m} \right| > \left| \left.\frac{d\hat{f}_{j,\mathrm{rRNA},i}}{dz_j}\right|_{z_j=z_m} \right| \tag{S97}$$

holds when $\left.\frac{d\hat{f}_{j,\mathrm{rRNA},i}}{dz_j}\right|_{z_j=z_m}$ and $\left.\frac{d\hat{f}_{j,\mathrm{rDNA},i}}{dz_j}\right|_{z_j=z_m}$ have opposite signs. Alternatively stated, when rRNA is increasing with an environmental variable while rDNA is simultaneously decreasing, then ratio rRNA:rDNA has a steeper response than rRNA alone, a differential response.

We assessed whether a given ASV disproportionately contributed to sensitivity results using leave-one-out removal. Specifically, each ASV was removed in turn from the set of $|S^*| = 21$ sensitivity differences $\{d_{j,i}\}$, and the Wilcoxon signed-rank test was re-run to the remaining $|S^*| - 1$ differences. No test was significant after BH-adjustment, indicating that no ASV disproportionately contributed to the sensitivity results.

### Figures

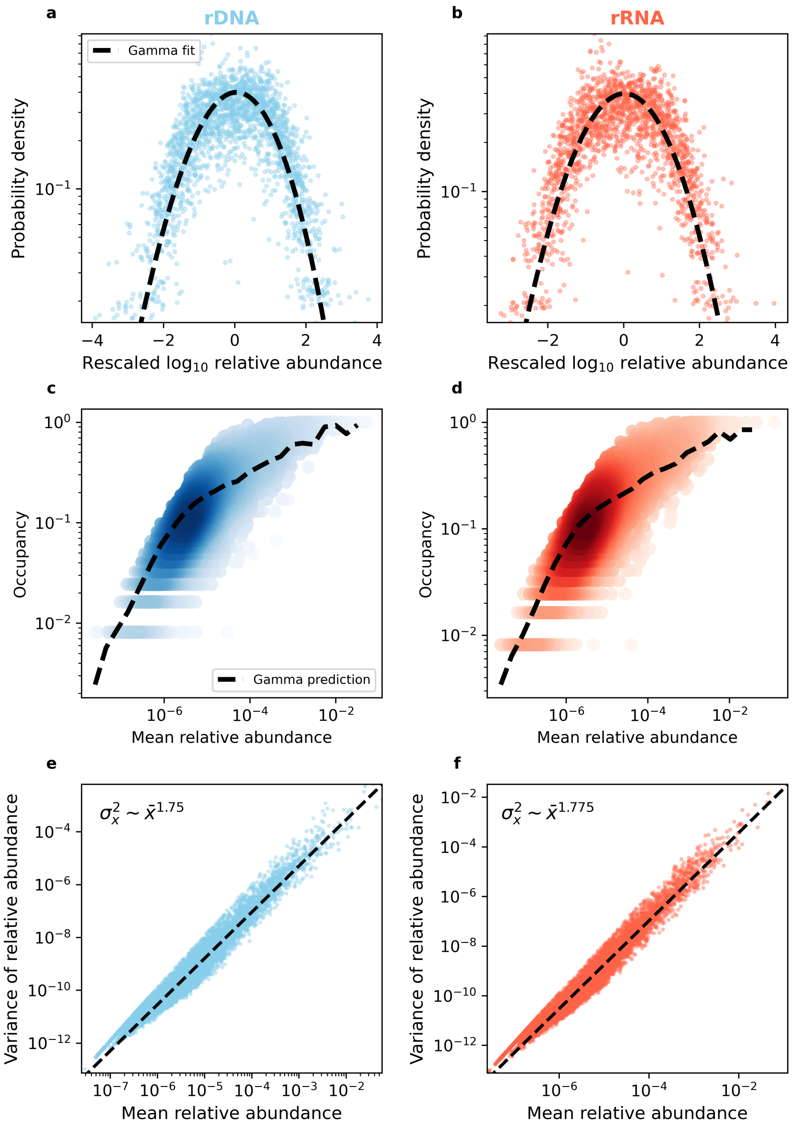


**Figure S1. rDNA and rRNA exhibit similar macroecological patterns.** Universal macroecological patterns including the **a,b)** gamma distributed Abundance Fluctuation Distribution (AFD), **c,d)** abundance-occupancy relationship, and **e,f)** Taylor's Law hold for both rDNA and rRNA.

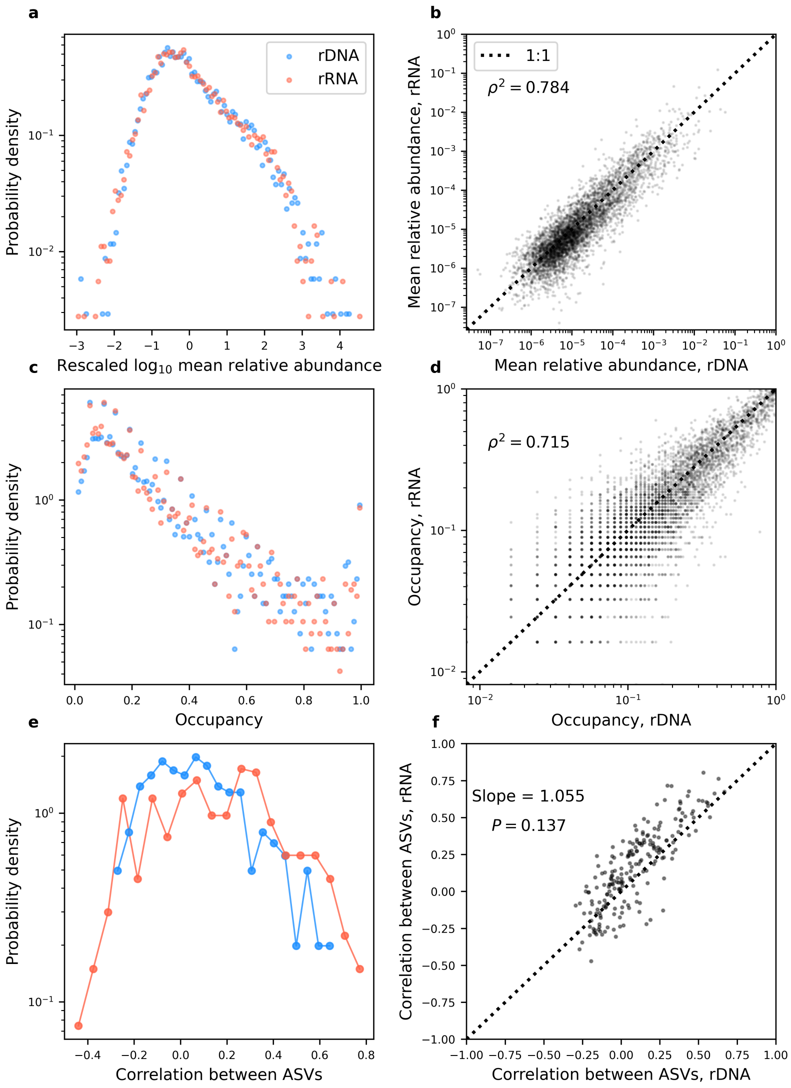


**Figure S2. Macroecological measures of abundance are highly correlated between rDNA and rRNA across ASVs. a,b)** Distributions of the mean relative abundance and occupancy are similar for both rDNA and rRNA as well as highly correlated across ASVs. **c)** Distributions of pairwise correlation coefficients between ASVs have slightly similar forms, but the deviation across ASVs is not statistically significant.

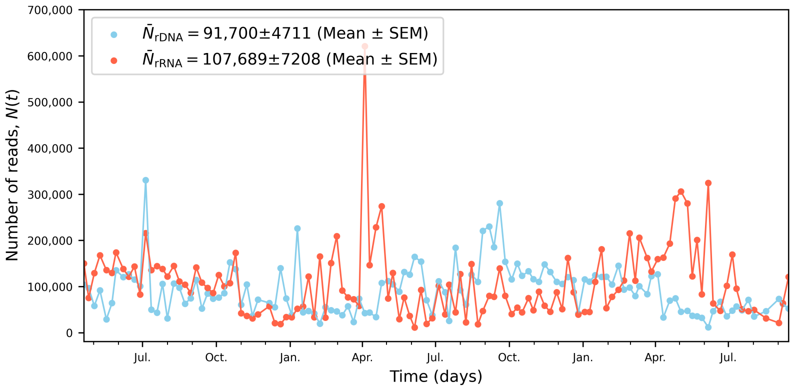


**Figure S3. Total number of reads over time for rDNA and rRNA** Consistently high numbers of reads allowed for 21 ASVs to be sampled at all timepoints even with abundances oscillating over four order of magnitude.

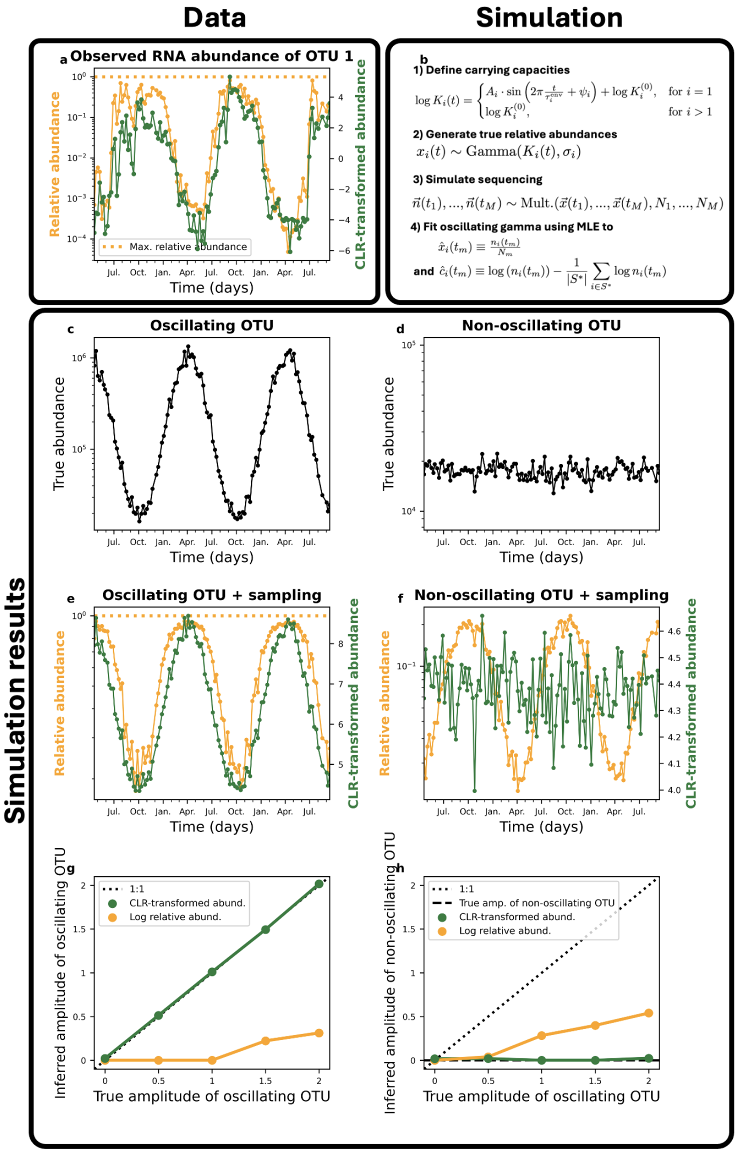


**Figure S4. Oscillations can be accurately inferred by considering compositionality. a)** In the data we identified a strongly oscillating ASV where the relative abundance began to saturate as it reached the maximum possible value of unity. In contrast, reads counts transformed using the log-ratio transformation (CLR) did not exhibit saturation, suggesting that the compositional nature of the data may shape observed oscillations. **b)** A minimal model that captures central patterns in the data was developed and simulations performed to assess the impact of compositionality (details in Materials and Methods). **c,d)** In the simulation, only one ASV had actual oscillatory dynamics while all remaining ASVs remained non-oscillatory. **e)** As expected, the saturating effect was observed using relative abundances but averted using CLR. **f)** The compositional nature of the data induced the appearance of strong oscillations in non-oscillating ASVs, while oscillatory artifacts were largely absent using CLR. **g)** The effect of compositionality had a severe effect on the ability to accurately infer the true amplitude of oscillations, while inference using CLR data was highly accurate. **h)** Similarly, inaccurate amplitudes were inferred for the non-oscillating ASV using relative abundances, while a near-zero amplitude was inferred using CLR.

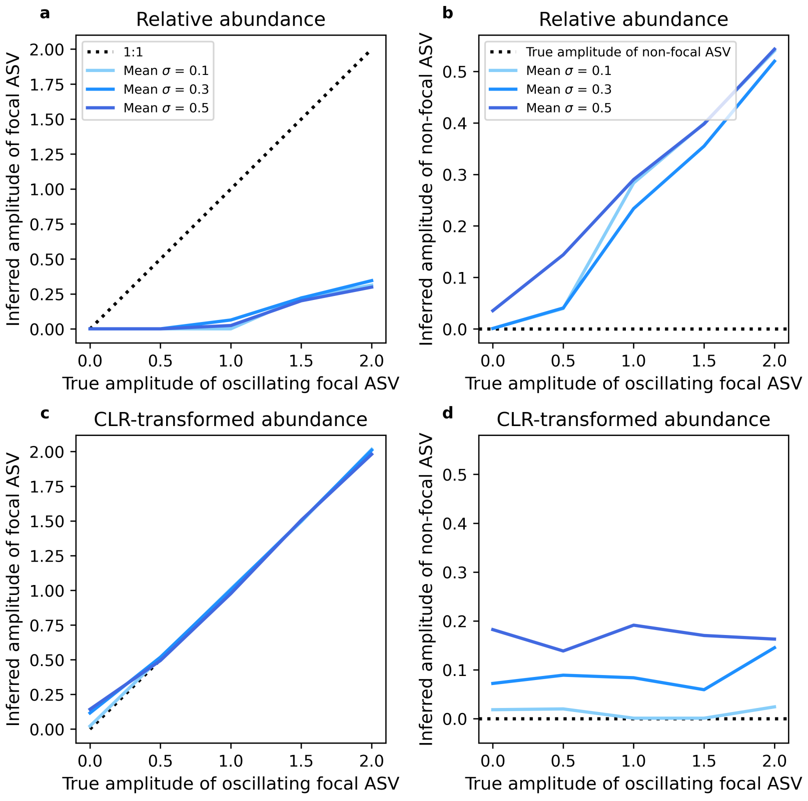


**Figure S5. Validity of amplitude inference for relative abundance vs. CLR transform.** The inference of the oscillatory amplitude $A_i$ was assessed for various strengths of environmental noise ($\sigma$). Amplitude inference was consistently incorrect for both **a,b)** oscillating and **c,d)** non-oscillating community members.

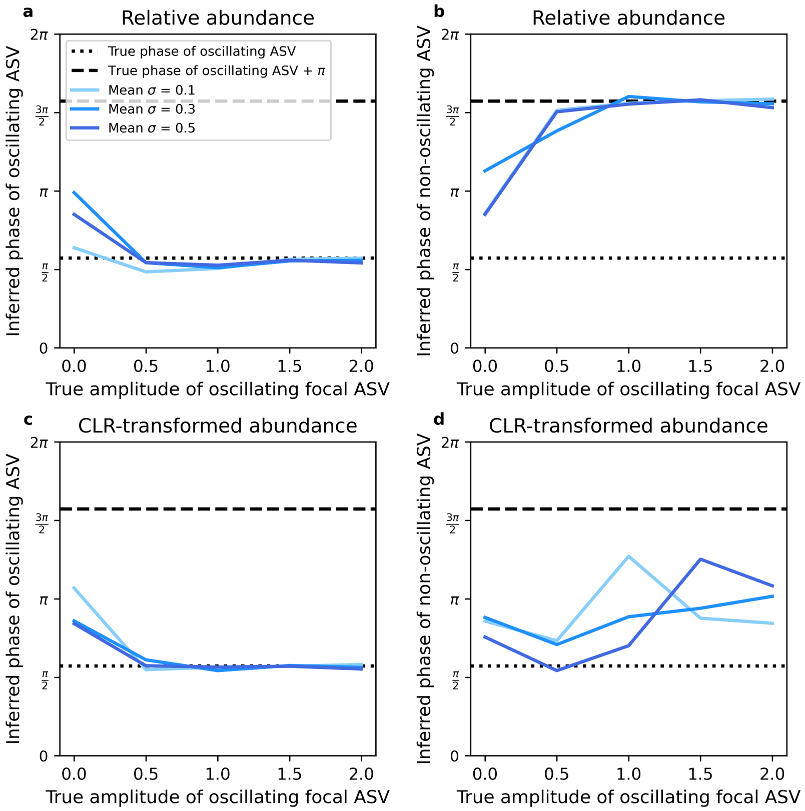


**Figure S6. Validity of phase inference for relative abundance vs. CLR transform.** Analogous analysis as Fig. S5 but for inference of the phase parameter $\psi_i$. **a,c)** The true phase of the oscillating community member could be accurately inferred for both relative abundance and CLR-transformed data. **b)** However, inference of the non-oscillating community member was heavily biased by the compositional nature of the data making it appear as if the dynamics oscillated with a shift of one unit $\pi$. **d)** This pattern was absent for CLR-transformed data.

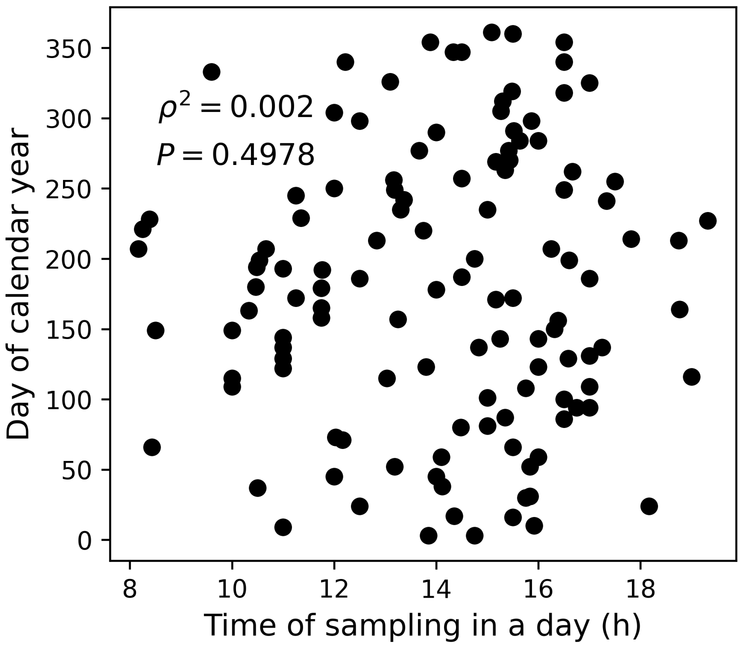


**Figure S7. Absence of bias in intra-daily sampling intervals.** There is no statistical relationship between the time of day in which a sample was acquired and its day within a year, implying that intra-daily variation in sampling time does not systematically bias our analysis.

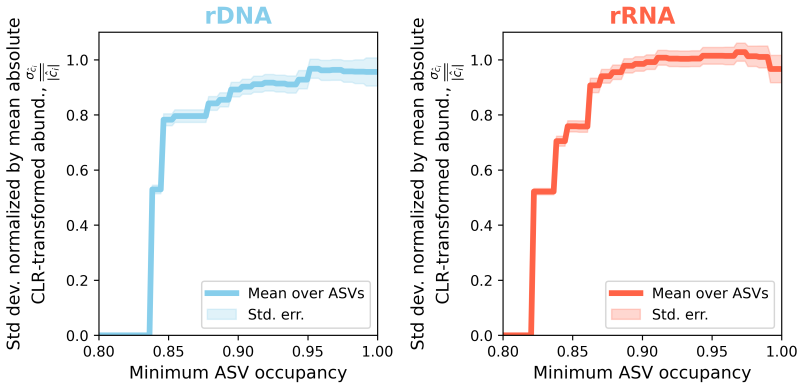


**Figure S8. Rarer ASVs do not systematically increase community-wide fluctuations.** ASVs following boom-bust growth dynamics can inflate fluctuations. To assess whether this influence was present in the data we identified sites where rarer ASVs were present and quantified the standard-deviation normalized by the mean of the absolute value (Supporting Information).

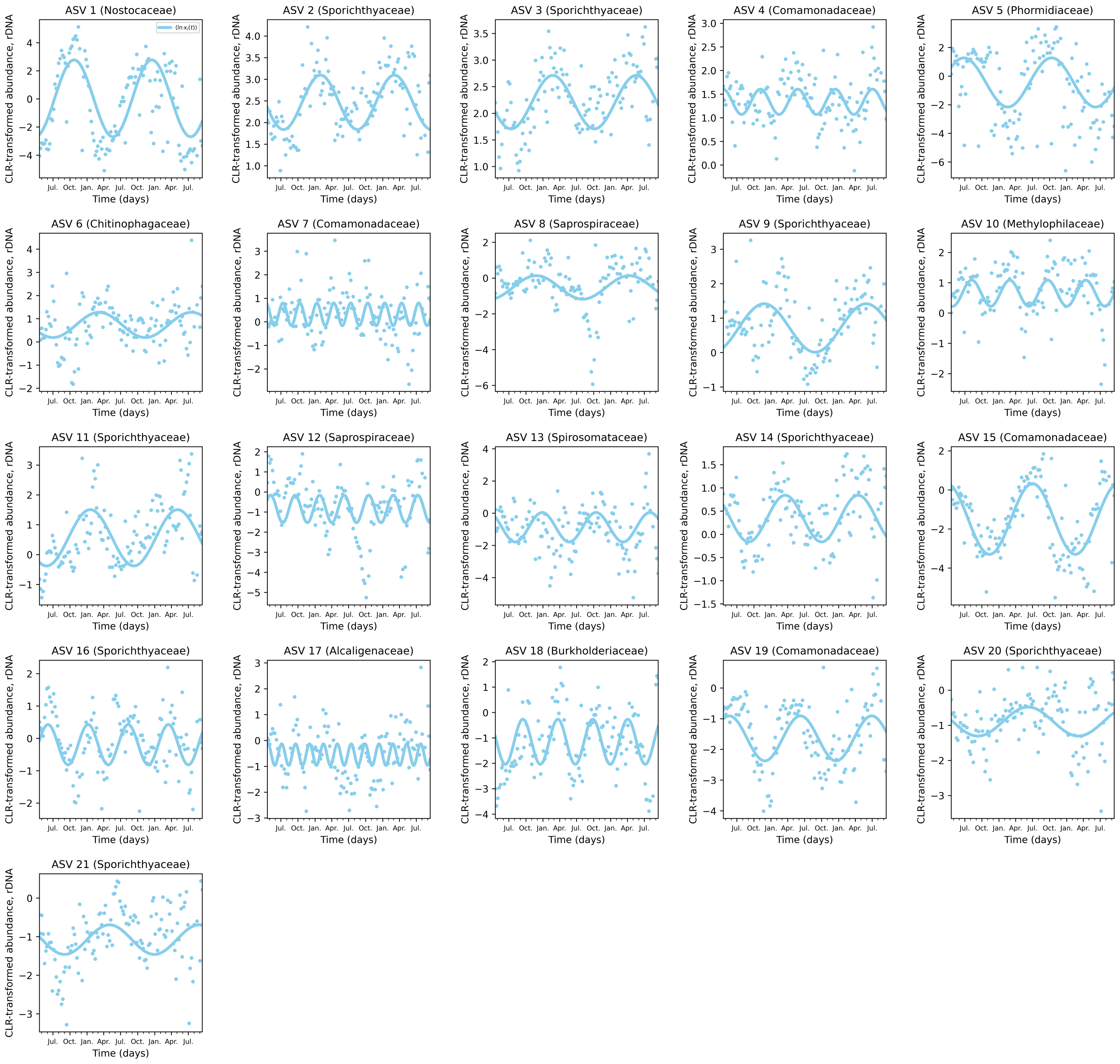


**Figure S9. Oscillatory fits for rDNA dynamics.** The model of an oscillating carrying capacity with gamma fluctuations provided reasonable fits for all ASVs present in every sample.

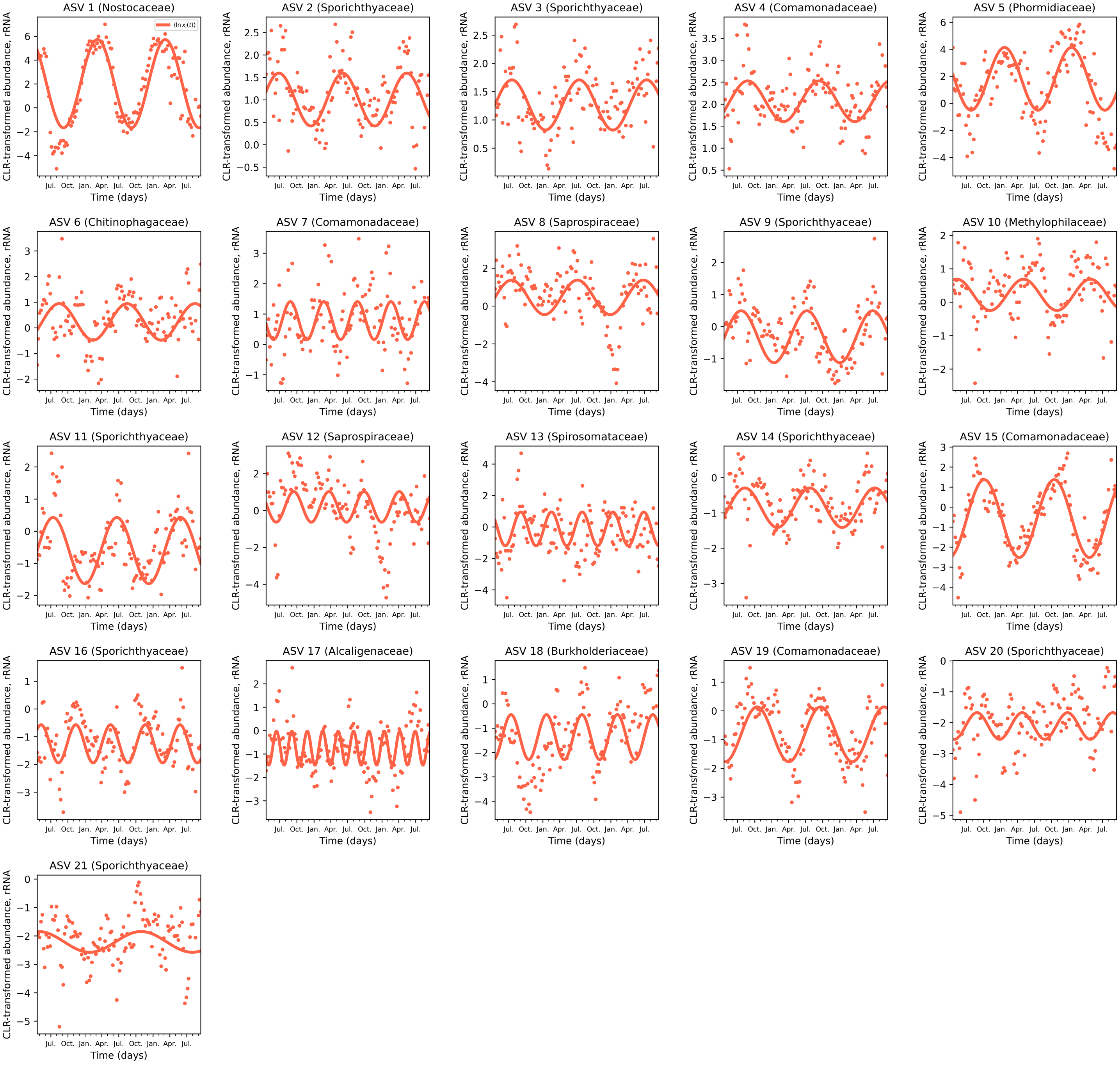


**Figure S10. Oscillatory fits for rRNA dynamics.** Equivalent plot as Fig. 3 for rRNA.

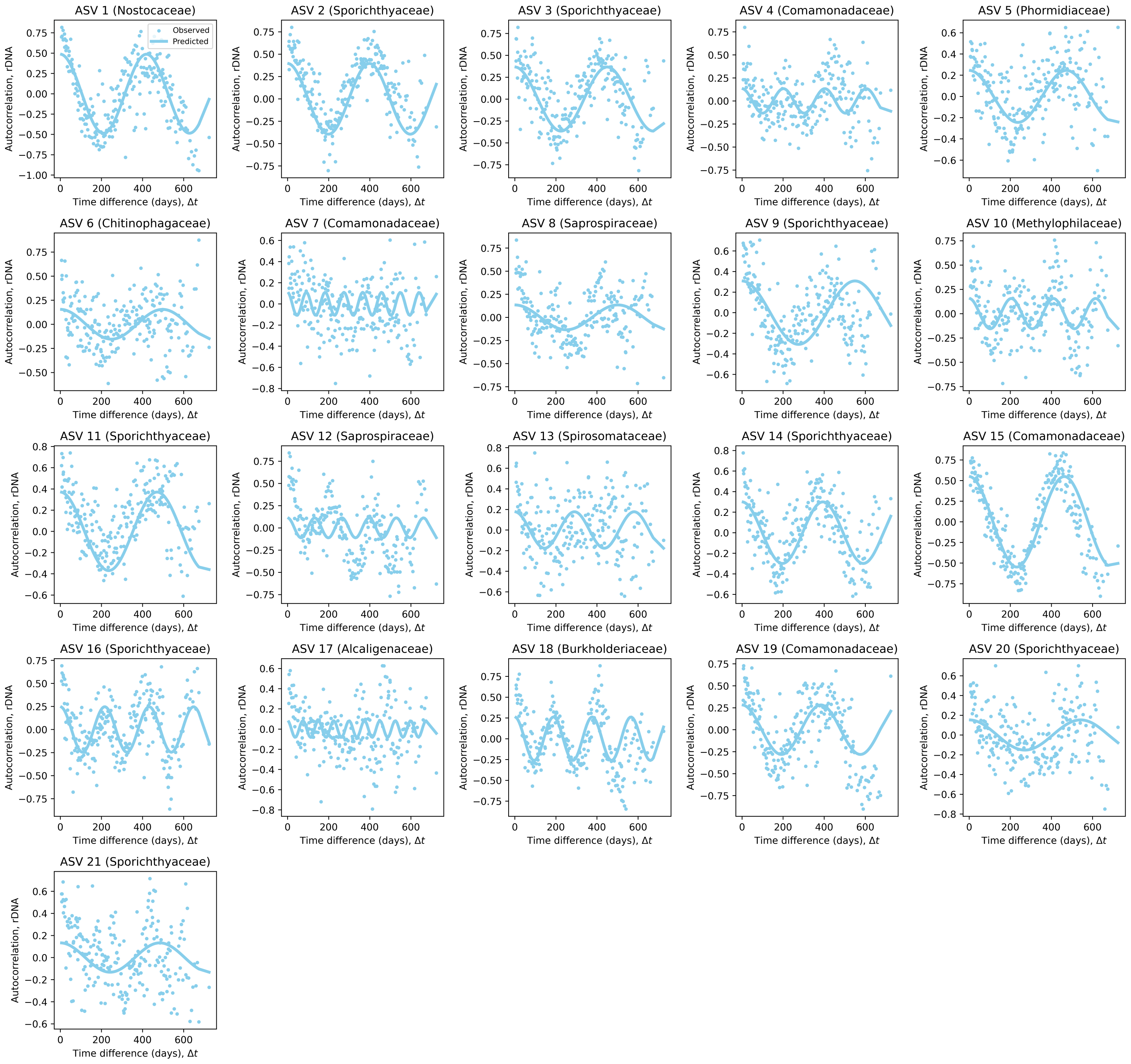


**Figure S11. Autocorrelation predictions for rDNA oscillations.** We derived a prediction of the autocorrelation function for the CLR-transformed abundance using the sine model (Eq. **??**). In general, the autocorrelation function reasonably captures empirical autocorrelation across ASVs.

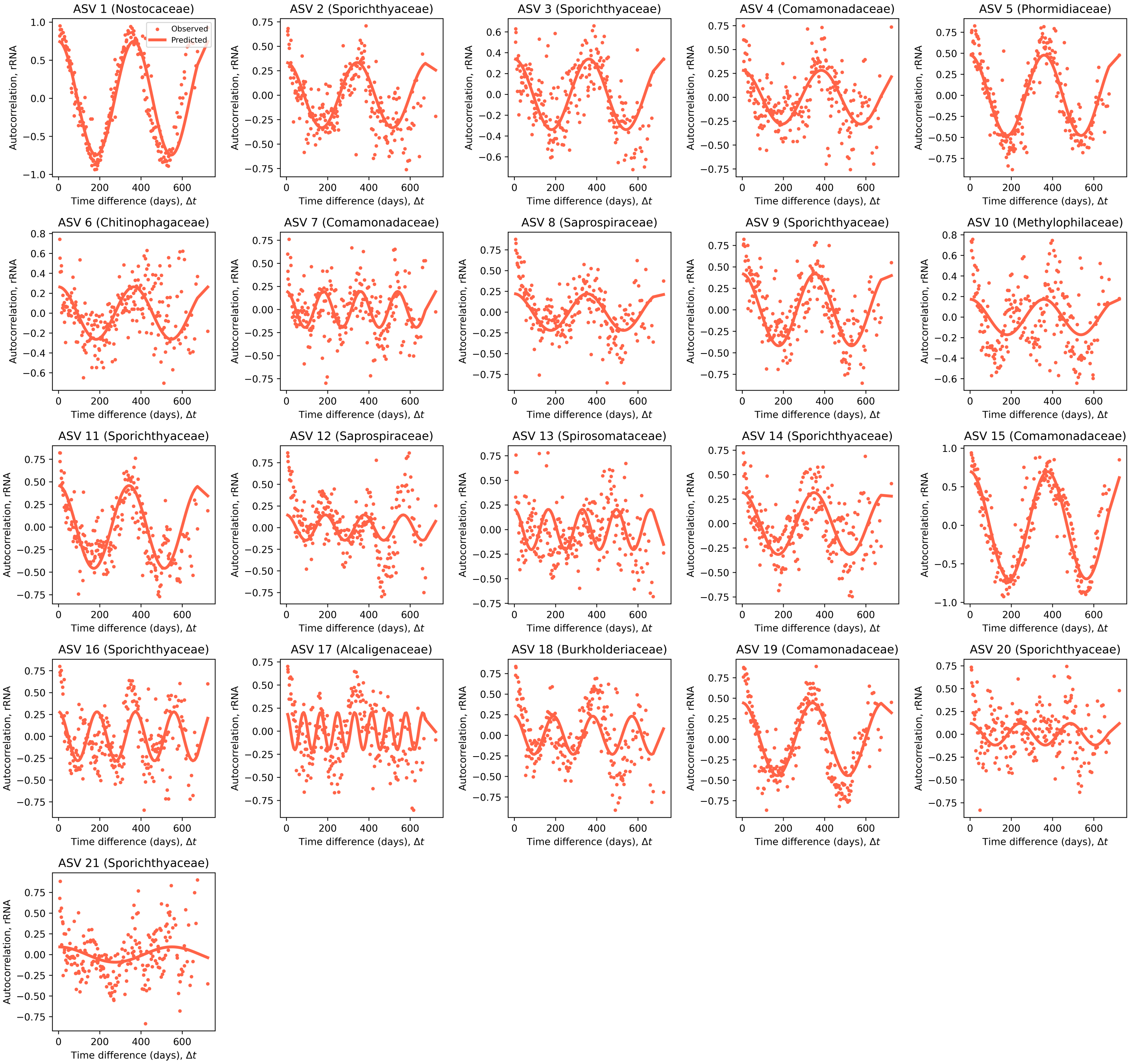


**Figure S12. Autocorrelation predictions for rRNA oscillations.** Equivalent plot as S11 for rRNA.

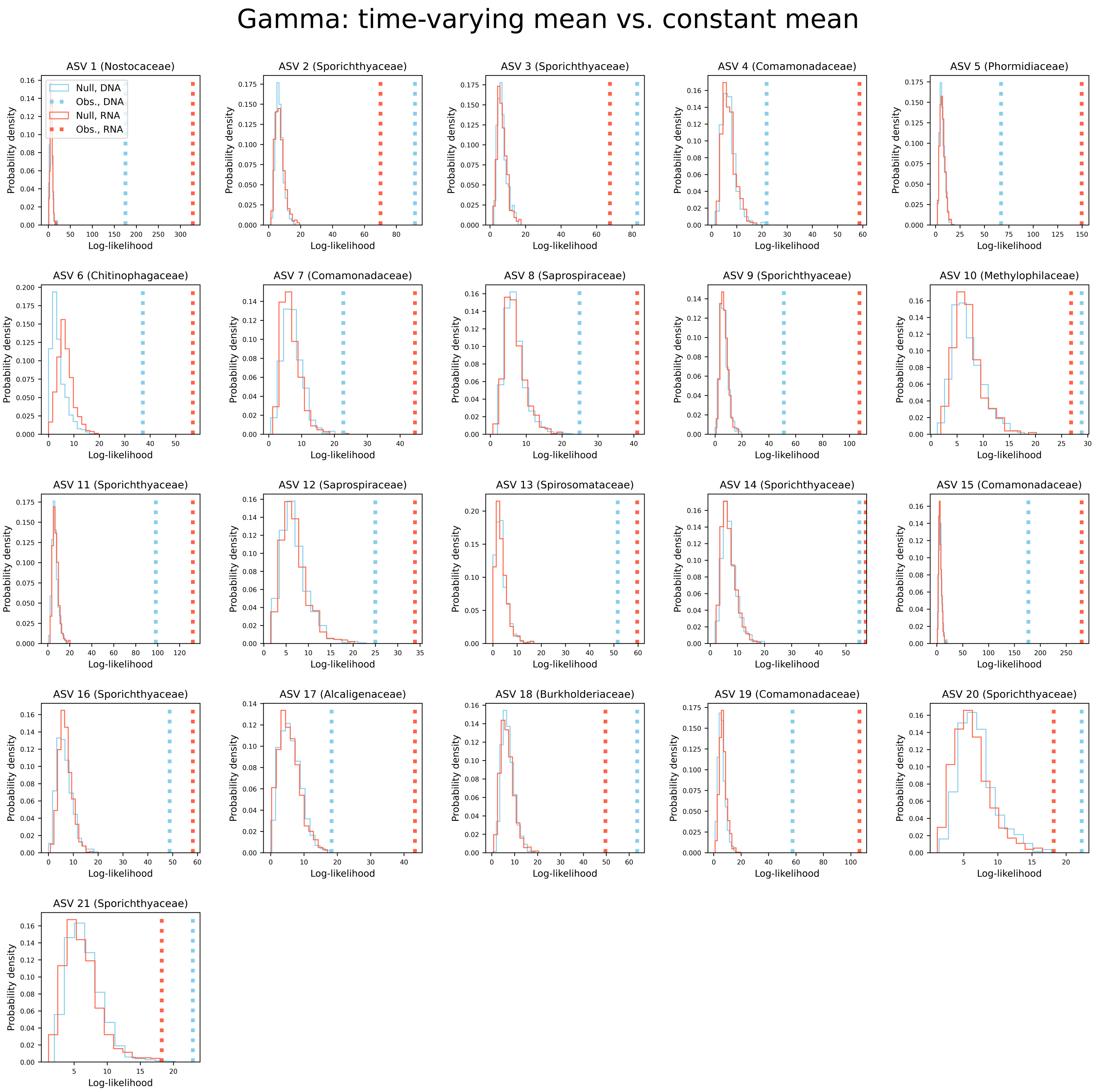


**Figure S13. Per-ASV Likelihood Ratio Test (LRT).** Bootstrapped LRTs performed per-ASV per data type.

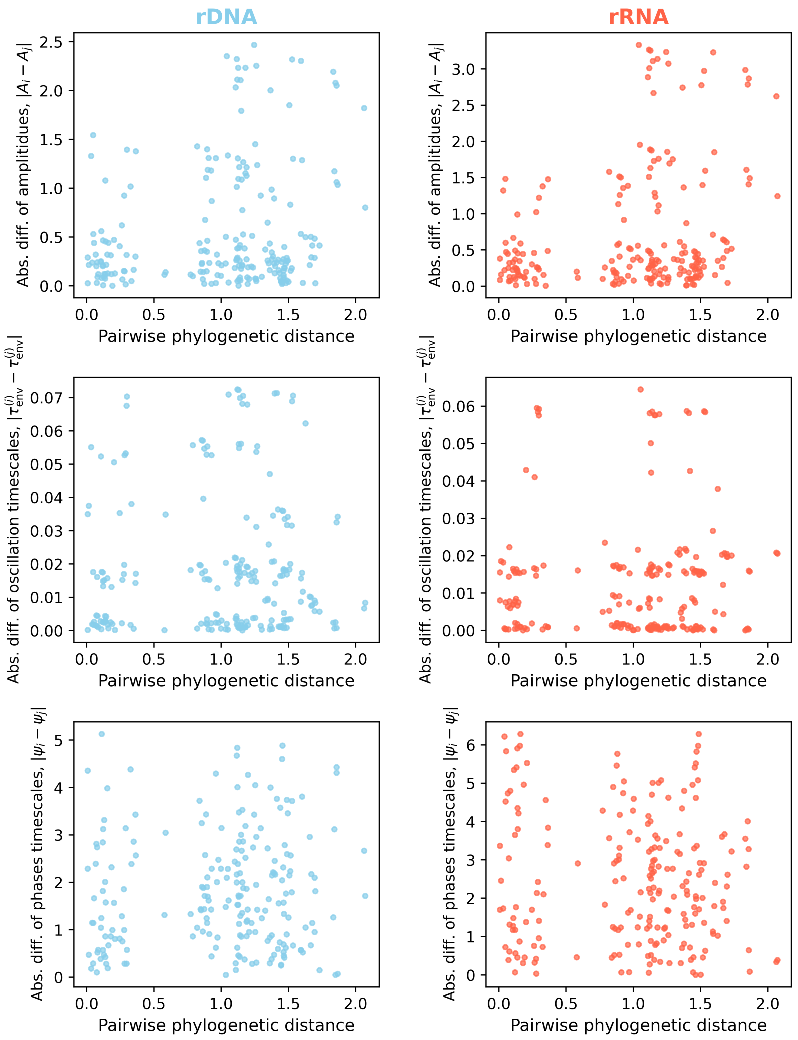


**Figure S14. Lack of a relationship between sine parameters and phylogenetic pairwise distance.** There is no apparent relationship between the pairwise phylogenetic distance between ASVs and the absolute difference in parameter values.

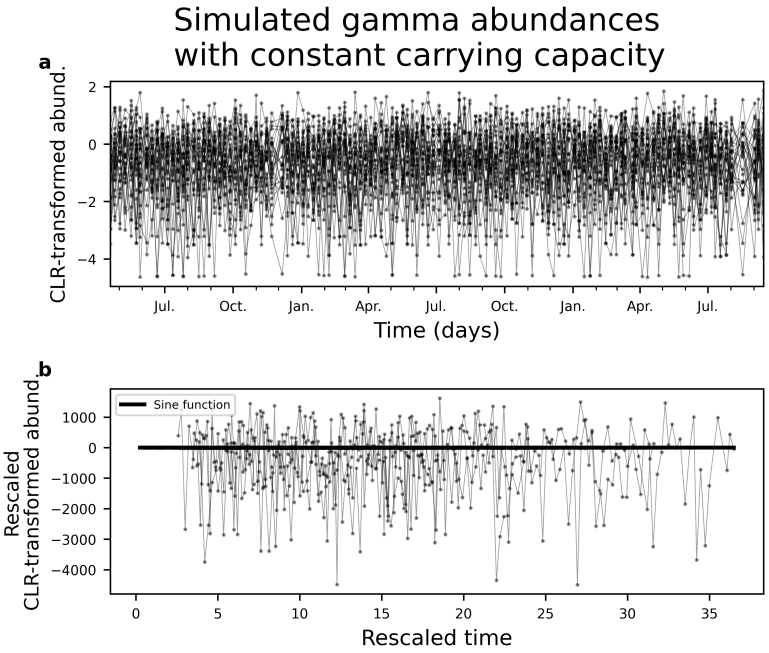


**Figure S15. The existence of a data collapse is not an artifact of statistical inference.** We did not observe oscillations when **a)** a sinusoidal function was fit to non-oscillating simulated data and **b)** the data was subsequently rescaled by the inferred parameters.

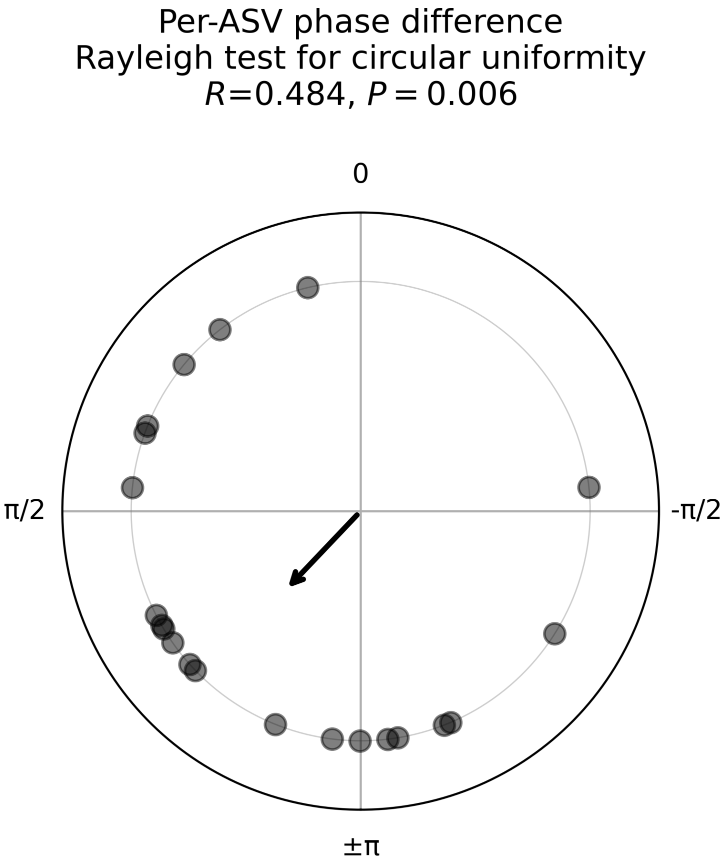


**Figure S16. ASV phase differences, $\Delta\psi_i$, are non-uniformly distributed on the unit circle.** Uniformity was tested by calculating the mean resultant length statistic across ASVs. The results are presented in polar coordinates.

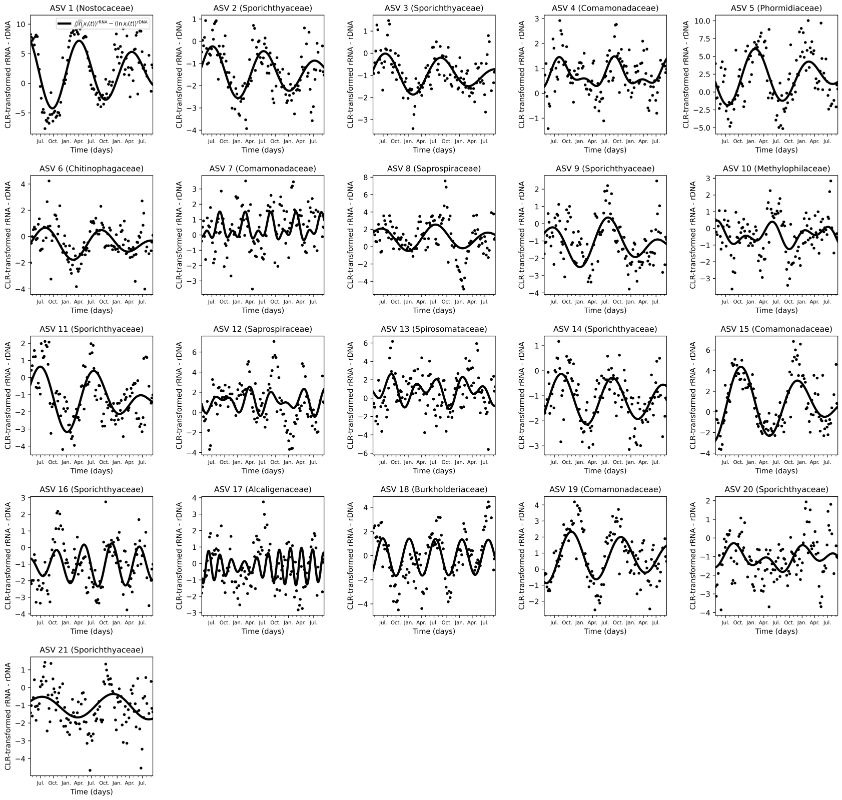


**Figure S17. rRNA:rDNA dynamics follow predictions from inferred oscillatory parameters.** Parameters inferred by fitting the model to rRNA and rDNA separately succeed in predicting rRNA:rDNA dynamics (Eq. **??**).

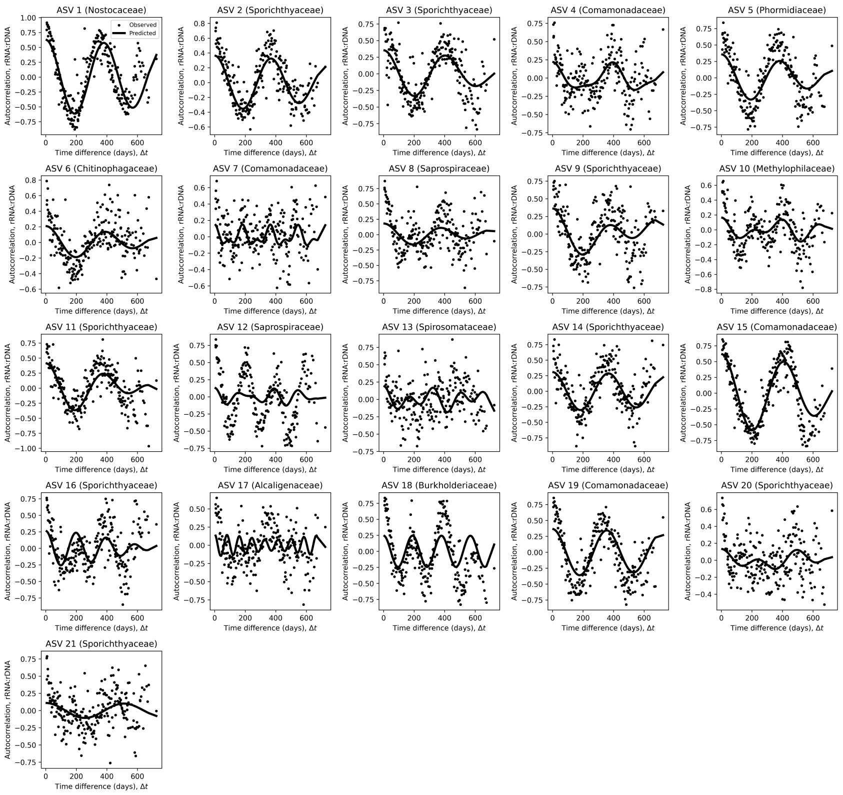


**Figure S18. Autocorrelation predictions for rRNA:rDNA.** We derived a prediction of the autocorrelation function for the CLR-transformed abundance of rRNA:rDNA using the sine model for rRNA and rDNA (Eq. **??**). The autocorrelation function of rRNA:rDNA reasonably captured empirical autocorrelation across ASVs using parameters fit separately to rRNA and rDNA.

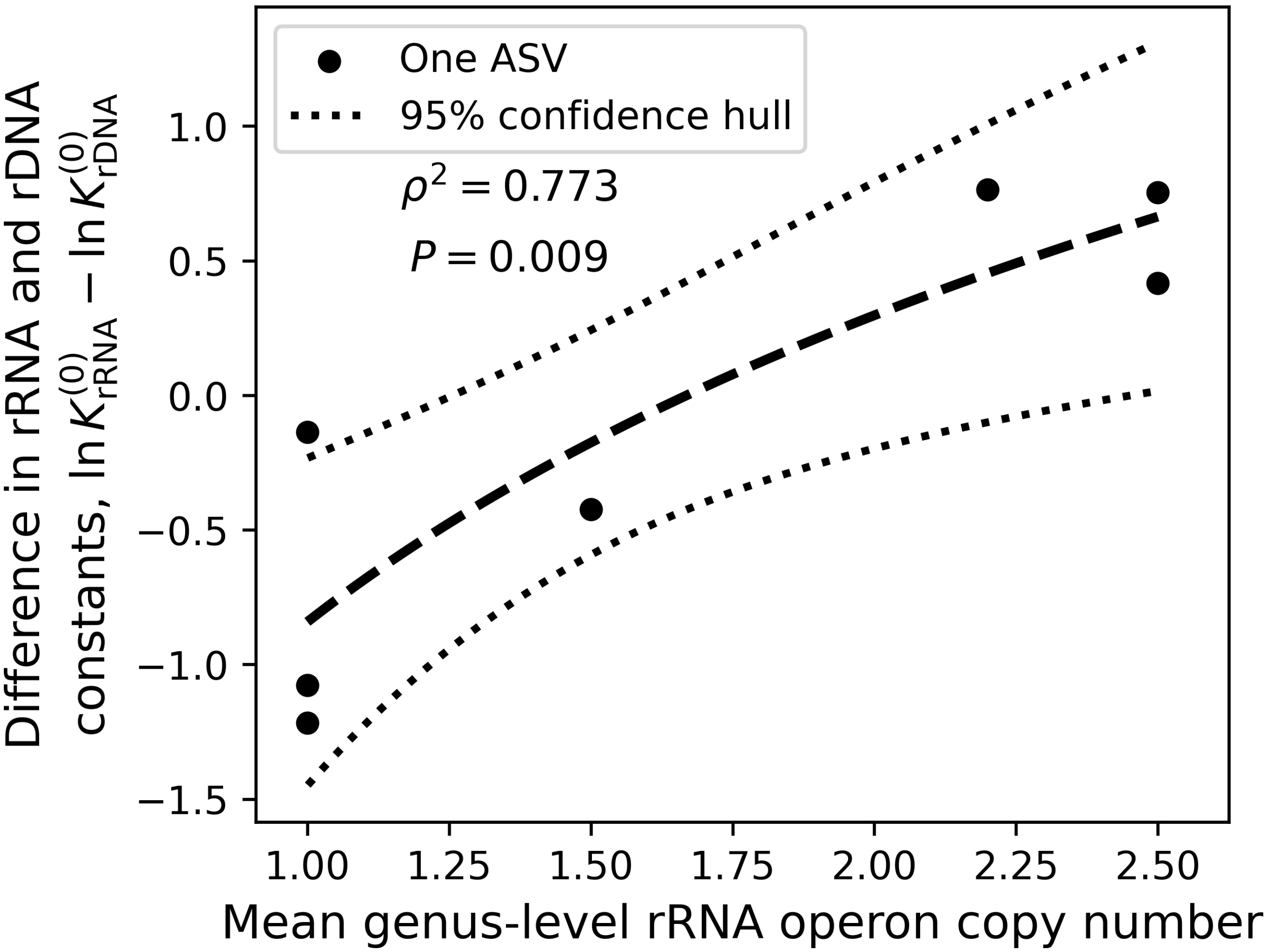


**Figure S19. Operon copy number vs. mean rRNA:rDNA relationship holds for inferred oscillatory parameter.** Analogous analysis of Fig. 4b for the intercept of the time-varying mean of our oscillating gamma model.

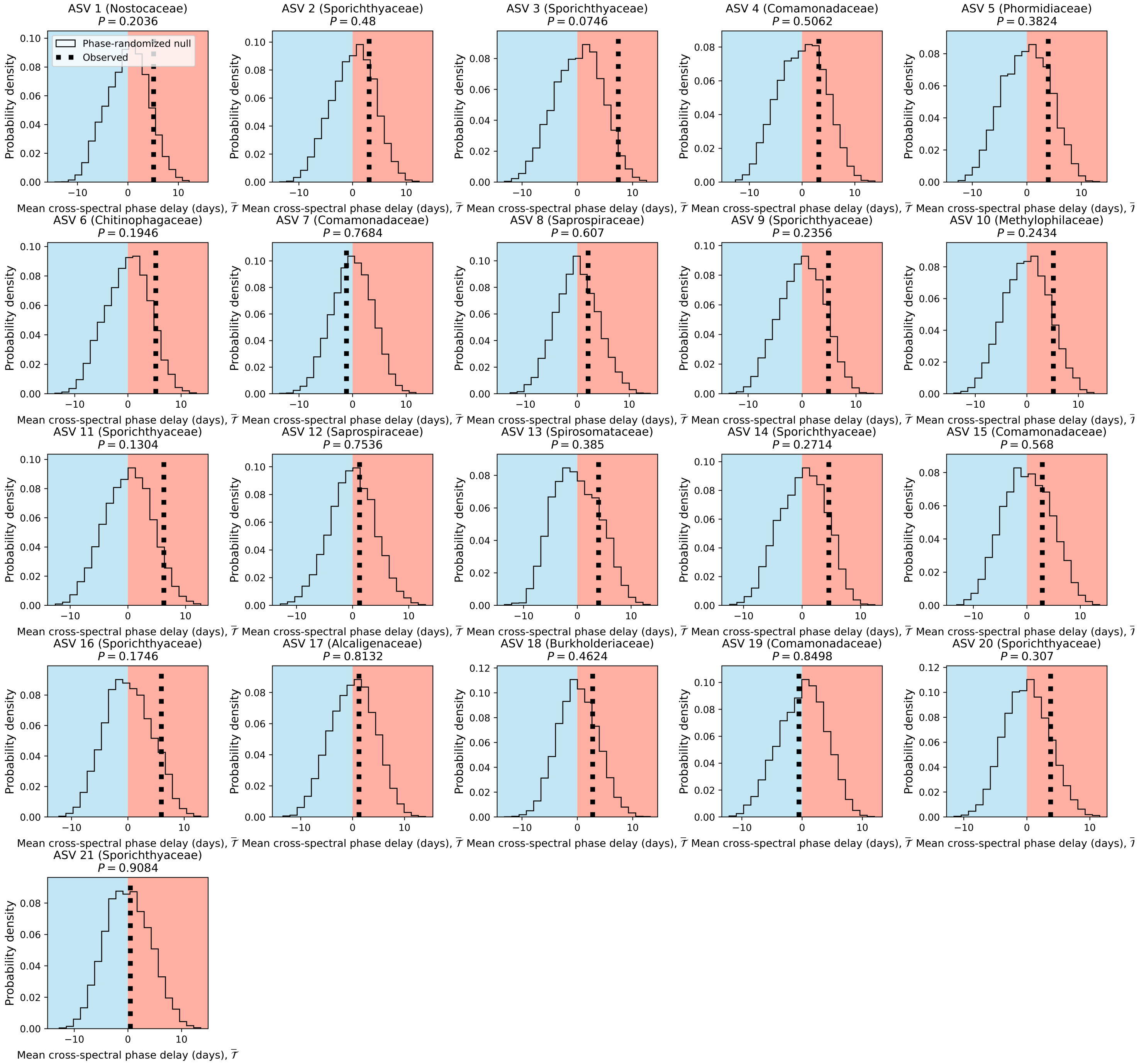


**Figure S20. Non-parametric analysis reveals the existence of weak, systematic time lag between rRNA and rDNA for *individual* ASVs.** Mean time lag between rRNA and rDNA were calculated using the Cross-Power Spectral Density and compared to a null where phases were drawn from a random distribution. Observed mean time lags were well-described by the bulk of the null distribution for all ASVs.

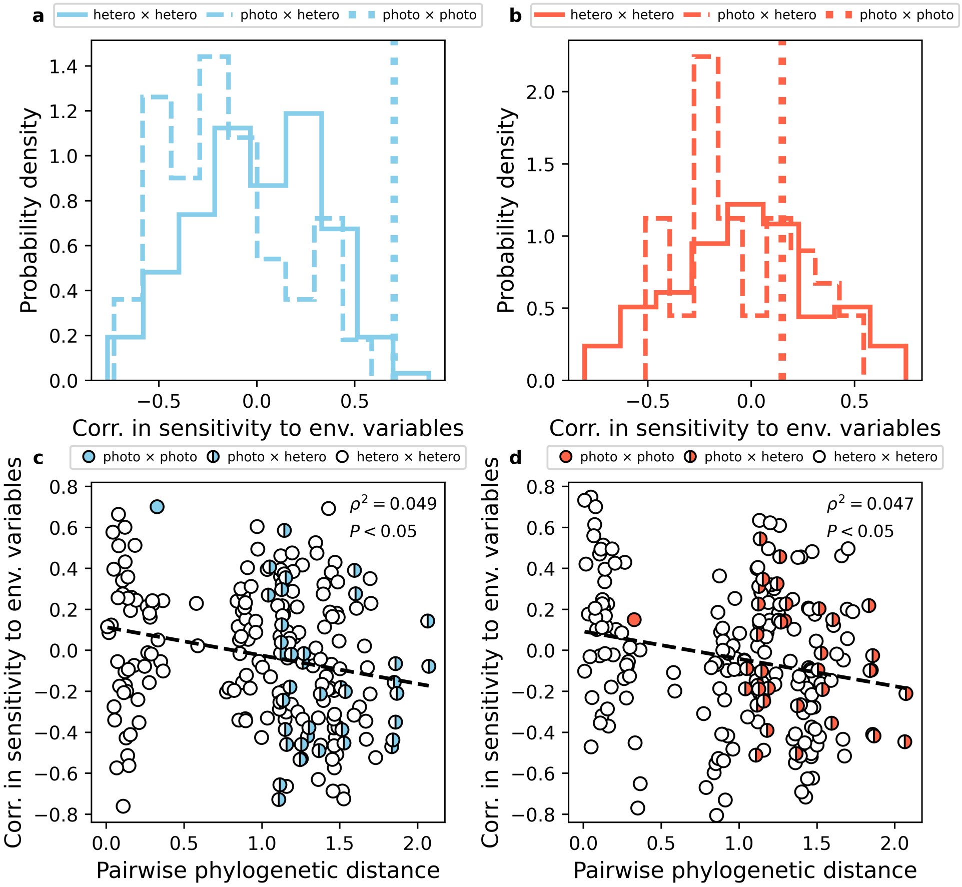


**Figure S21. ASV sensitivities to environmental variables vary by trophic level, phylogeny, and data type. a)** A strong positive correlation was found between the sole phototroph × phototroph comparison in rDNA that was absent for rRNA. **b)** Significant phylogenetic distance-decay relationships were found in both rDNA and rRNA.
